# Single-cell-resolved calcium and organelle dynamics in resistosome-mediated cell death

**DOI:** 10.1101/2025.06.27.662017

**Authors:** Yi-Feng Chen, Kuan-Yu Lin, Ching-Yi Huang, Liang-Yu Hou, Enoch Lok Him Yuen, Wei-Che Sun, Bing-Jen Chiang, Chin-Wen Chang, Hung-Yu Wang, Tolga O. Bozkurt, Chih-Hang Wu

## Abstract

Plant nucleotide-binding domain leucine-rich repeat-containing (NLR) proteins act as intracellular immune receptors that assemble into resistosomes to execute immune responses. However, the subcellular processes during cell death following resistosome activation remain unclear. Here, we visualized the changes in calcium signaling and organelle behavior after activation of the NRC4 (NLR-required for cell death 4) resistosome. We found that NRC4 membrane enrichment coincided with calcium influx. This is followed by sequential mitochondria and plastid disruption, endoplasmic reticulum fragmentation and cytoskeleton depolymerization. Subsequent loss of plasma membrane integrity, nuclear shrinkage, and vacuolar collapse mark the terminal stage of cell death. Our findings reveal a spatiotemporally-resolved cascade of subcellular events downstream of resistosome activation, providing new mechanistic insight into the execution phase of plant immune cell death.

## Introduction

Nucleotide-binding domain leucine-rich repeat receptors (NLRs) play critical roles in the innate immunity of both plants and animals (Duxbury et al. 2021). In animals, activated NLRs form inflammasomes that guide Gasdermin oligomers to the plasma membrane, resulting in pyroptotic cell death and the elimination of pathogens. In plants, activated NLRs form oligomeric structures known as resistosomes (Duxbury et al. 2021; Chai et al. 2023). These resistosomes can function as enzymes that generate small secondary messenger molecules or as calcium channels targeting the plasma membrane, ultimately triggering hypersensitive cell death to restrict pathogen invasion (Chai et al. 2023).

The NLR required for cell death (NRC) family comprises multiple helper NLRs with partial functional redundancy, acting downstream of several sensor NLRs (Wu et al. 2017). For example, the *Nicotiana benthamiana* helper NLR NRC4 works with both the Potato virus X (PVX) resistance protein Rx and the late blight resistance protein Rpi-blb2, whereas NRC2 functions with Rx but not Rpi-blb2. Upon activation, NRCs form high molecular weight resistosome complexes and localize at the plasma membrane as puncta (Duggan et al. 2021; Ahn et al. 2023; Contreras et al. 2023). Recent cryogenic electron microscopy (cryo-EM) studies showed that NRC2 and NRC4 exist as autoinhibited dimers at the resting state (Ma et al. 2024; Selvaraj et al. 2024). Upon activation by sensor NLRs or autoactivation mutations, NRCs oligomerize into hexameric resistosome complexes (Liu et al. 2024; Madhuprakash et al. 2024). Activated NRCs likely function as calcium channels, triggering a calcium influx into the cytosol and initiating downstream immune responses (Liu et al. 2024). However, how resistosome activation and calcium signaling drive subcellular reorganization and culminate in cell death remains poorly understood.

A major obstacle in dissecting these events has been the rapid onset of hypersensitive cell death following resistosome activation. Consequently, most studies have relied on death-deficient NLR variants or immune-compromised mutant backgrounds, limiting the ability to monitor dynamic cellular changes during the execution phase. To overcome this, we employed a recently developed copper-inducible transient expression system in *N. benthamiana* (Chiang et al. 2024), allowing us to temporally control resistosome formation and perform live-cell imaging to uncover the subcellular events that accompany NRC4-mediated cell death.

## Results

### NRC4 forms the resistosome complex and triggers cell death within three hours of copper-induced effector expression

To resolve the timing of cell death under the copper-inducible system, we established a leaf disc-based cell death assay using autofluorescence as a readout. We transiently expressed Rpi-blb2 along with either inducible AVRblb2 from *Phytophthora infestans* or inducible CP from PVX in *N. benthamiana* leaves. Two days post-agroinfiltration, we infiltrated copper solution into the leaves and punched leaf discs for real-time fluorescence measurement in a plate reader (Fig. S1A). Autofluorescence in samples expressing copper-inducible AVRblb2 rose immediately and saturated ∼3 h post-copper infiltration (hpci), indicating that Rpi-blb2 triggers cell death efficiently within this window (Fig. 1A). By contrast, samples expressing negative control, the copper-inducible CP, did not show any considerable increase in fluorescence. The AVRblb2-induced autofluorescence signal was abolished in the *nrc2/3/4* triple knockout (*nrc*) background but was restored by expression of wild-type NRC4 (Fig. 1B), whereas previously characterized cell death deficient NRC4^L9E^, NRC4^K190R^ mutants or NRC2 failed to complement the phenotype (Fig. 1B). These results are consistent with the visible cell death phenotype observed at 2 days post copper infiltration (dpci) (Fig. S1B and S1C).

**Fig. 1.**
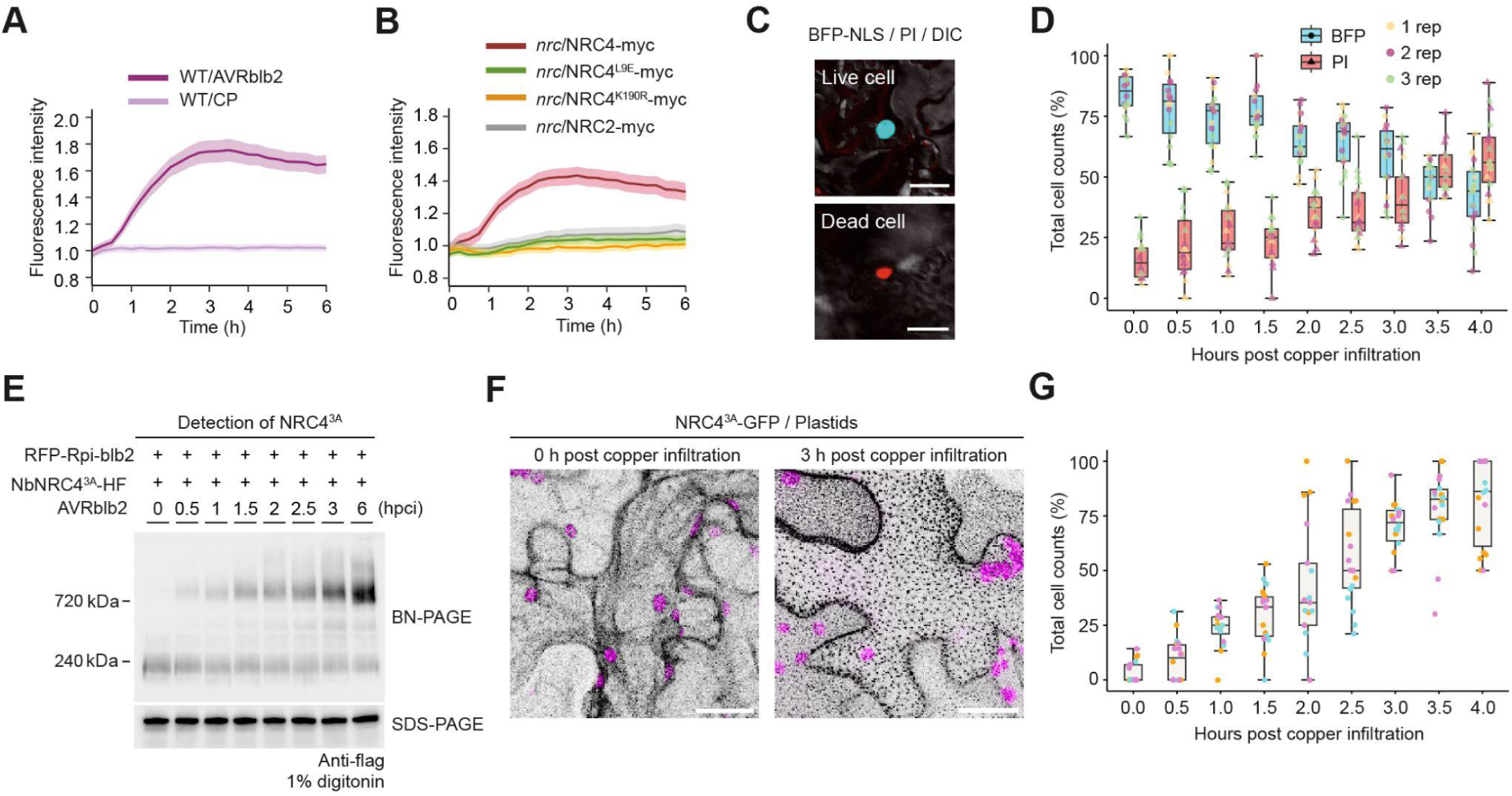
Copper-inducible effector expression triggers rapid hypersensitive cell death and NRC4 resistosome formation in *N. benthamiana*. A. Quantification of cell death in a leaf disc-based assay upon copper-inducible effector expression. Leaves were co-infiltrated with *35S::Rpi-blb2* and a copper-inducible effector construct (*AVRblb2* or *CP*). Autofluorescence was measured using a plate reader following copper infiltration. Fluorescence intensities were normalized to mock-treated controls. Solid lines represent mean values; shaded areas indicate standard error (n = 34–36 discs from 3 independent experiments). B. Complementation assay in *nrc2/3/4* triple knockout (*nrc*) *N. benthamiana* plants. NRC4, NRC4 mutants (L9E, K190R), or NRC2 were expressed along with Rpi-blb2 and inducible AVRblb2. Autofluorescence was measured as in (A). C. Representative confocal images showing a live cell (top) with nuclear BFP (BFP-NLS) and a dead cell (bottom) stained with propidium iodide (PI). Scale bar = 30 μm. D. Time-course quantification of live (BFP-NLS-positive) and dead (PI-positive) cells following copper treatment. Box plots show the percentage of total cells that are either live or dead at each time point. Data represent three independent biological replicates (n = 18 images). E. Time-course analysis of NRC4^3A^-HF resistosome assembly. Blue native PAGE (BN-PAGE) shows the accumulation of high molecular weight NRC4^3A^-HF complexes after AVRblb2/RFP-Rpi-blb2 activation at the indicated hours post copper infiltration (hpci). SDS-PAGE was used to assess total NRC4^3A^-HF protein as a loading control. F. Confocal Z-stack maximum projection showing subcellular localization of NRC4^3A^-GFP at 0 and 3 hpci. Gray: NRC4^3A^-GFP; magenta: plastid autofluorescence. Leaves were co-expressed with *35S::NRC4^3A^-GFP*, *35S::Rpi-blb2*, and *CBS4::AVRblb2* in the *nrc* background. Images were taken 2 days post infiltration. Scale bar = 20 μm. G. Time-course quantification of NRC4^3A^-GFP puncta formation. Box plots show the percentage of cells with visible NRC4^3A^-GFP puncta at each time point post copper infiltration. Data represent three independent biological replicates (n=16-19 images).

To assess cell viability under a confocal microscope, we transiently expressed a blue fluorescent protein (mTagBFP2) fused to nuclear localization signal (NLS) and performed propidium iodide (PI) staining. Time-course experiments showed that the percentage of epidermal cells with PI-positive nuclei (indicating dead cells) increased over time, while the percentage of cells displaying nuclear BFP signal (indicating live cells) decreased correspondingly (Fig. 1C and 1D). To determine whether NRC4 high molecular weight resistosome complexes form within this timeframe, we conducted time-course native-PAGE assays using the NRC4^L9A/V10A/L14A^ variant (hereafter referred to as NRC4^3A^) (Wang et al. 2025). NRC4^3A^ resistosome high molecular weight complexes became detectable as early as 0.5 hours post-induction and continued to accumulate at 3 and 6 hours (Fig. 1E). Additionally, we observed that activated NRC4^3A^-GFP formed puncta on the plasma membrane (Fig. 1F). A time-course analysis revealed that the proportion of cells exhibiting NRC4^3A^ puncta increased over time, reaching approximately 80% by 3 hours post-AVRblb2 induction (Fig. 1G). Collectively, these findings demonstrate that Rpi-blb2/AVRblb2-triggered, NRC4-dependent cell death can be efficiently and reproducibly activated using the copper-inducible system.

### Activated wild-type NRC4 forms puncta and filamentous structures prior to cell collapse

Next, we used the inducible system to investigate the subcellular dynamics of NRC4 activation using time-lapse imaging. First, we compared the dynamics of NRC4^3A^ and NRC4 upon activation by Rpi-blb2 and AVRblb2. We transiently expressed NRC4-GFP variants, untagged Rpi-blb2, and inducible AVRblb2 in *N. benthamiana* leaves. Two days after agroinfiltration, we infiltrated copper solution into the leaves to induce effector expression. Leaf discs were then immediately prepared for confocal microscopy to observe NRC4 dynamics (Fig. S1A). Since NRC4 forms puncta at the plasma membrane, we adjusted the focal plane to the cell periphery for time-lapse imaging. At the early stage of time-lapse imaging, NRC4^3A^-GFP displayed highly dynamic behavior reflecting its diffused cytosolic distribution at the resting state. Around 30 minutes to an hour (timing varied between cells) post induction, small punctate structures began to appear. These puncta were immobile and gradually increased in intensity over time, likely reflecting the clustering of resistosome complexes at the plasma membrane (Fig. 2A, 2C, S2A, S3A; Movie S1). Experiments using cytosolic mRFP revealed that the cytosol remains dynamic following the formation of NRC4^3A^-GFP puncta (Movie S2), with reduced GFP/mRFP co-localization coefficient value post-NRC4 puncta formation (Fig. S3B). Fluorescence recovery after photobleaching (FRAP) assays confirmed that these NRC4^3A^ puncta are not mobile (Fig. S3C-D).

**Fig. 2.**
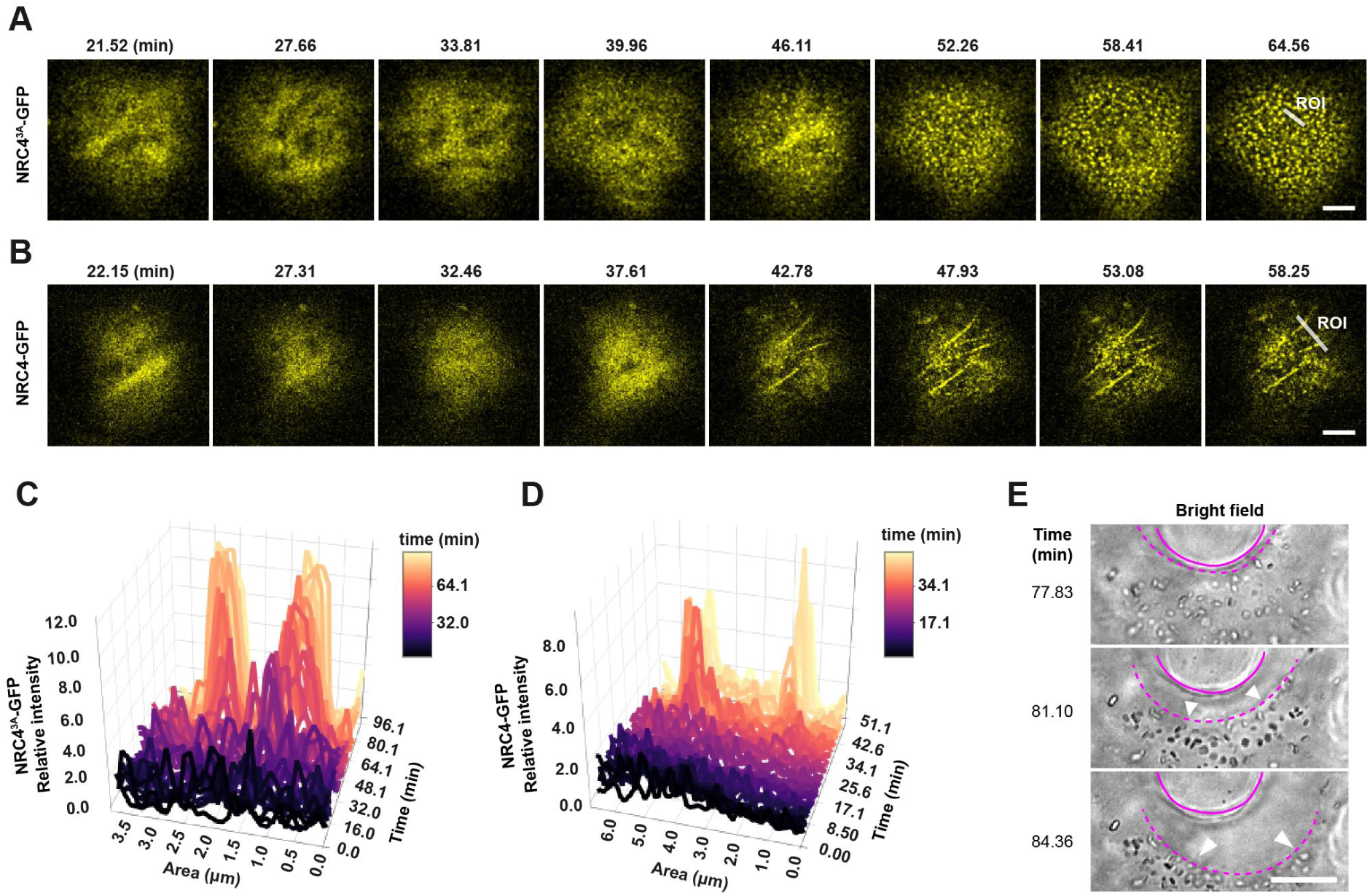
NRC4 forms resistosome puncta and filamentous structures prior to cell collapse. A. and B. Time-lapse confocal images showing subcellular localization dynamics of (A) NRC4^3A^-GFP and (B) NRC4-GFP following copper-induced expression of AVRblb2. *N. benthamiana nrc* plants were co-infiltrated with *35S::Rpi-blb2*, *CBS4::AVRblb2*, and *35S::CUP2-p65*. At 2 days post infiltration, copper solution was applied to induce effector expression, followed by confocal imaging at the cell periphery. White lines in the final panels indicate regions of interest (ROIs) used for intensity profiling in (C) and (D). Scale bar = 5 μm. (See Movies S1 and S3.) C. and D. Time-course fluorescence intensity profiles of (C) NRC4^3A^-GFP and (D) NRC4-GFP along the ROIs marked in (A) and (B), respectively. Each line represents fluorescence intensity across the ROI at the indicated time point. A pseudocolor gradient reflects the progression of time. E. Bright-field time-lapse images showing morphological changes during NRC4-GFP–induced cell collapse. Solid and dotted lines outline the upper and lower cell boundaries, respectively. White arrowheads indicate vacuolar shrinkage. Scale bar = 10 μm. (See Movie S3.)

We then conducted time-lapse imaging with NRC4-GFP. Similar to NRC4^3A^, NRC4 exhibits dynamic cytosolic movement initially. However, unlike NRC4^3A^, NRC4-GFP abruptly formed puncta along with filament-like structures (Fig. 2B, 2D, S2B; Movie S3). These puncta and filaments remained immobile until the cell collapsed (Fig. 2D; Movie S3). Cytoplasmic streaming ceased at the onset of puncta/filament formation (Movie S4), coinciding with a sharp drop in the co-localization coefficient value between NRC4-GFP and cytosolic mRFP (Fig. S2E). Within 15–30 minutes following the appearance of these structures, the plasma membrane detached from the cell wall, overall fluorescence weakened, and the NRC4–GFP signal disappeared (Fig. S2B, 2E; Movie S3).

### Cytoplasmic calcium influx coincides with NRC4 membrane enrichment

Previous studies have shown that NRC4 activation induces calcium signaling in leaves (Liu et al. 2024), but the kinetics of this influx at single-cell resolution have remained unclear. To address this, we generated a stable transgenic *N. benthamiana* line expressing the Ca²⁺ reporter GCaMP6 and performed copper-inducible cell-death assays described above, imaging calcium dynamics in individual cells (Fig. S1A) (Chen et al. 2013). Roughly 30–60 min after copper induction (varying between cells), GCaMP6 fluorescence rose sharply, indicating a rapid increase in cytosolic and nuclear Ca²⁺ (Fig. 3A-B, S4A, S5A; Movie S5). The calcium signal peaked within 3-5 minutes of influx onset and returned to baseline over the next 3-5 minutes (Fig. 3A-B, S5A, and Fig. S5B). Then, rapid cell collapse occurred, marked by the detachment of the plasma membrane from the cell wall at approximately 15 minutes or longer after the calcium peak (Fig. 3A-B; Movie S5). Thus, we conclude that NRC4 activation elicits a transient Ca²⁺ influx that precedes the onset of cell death. To confirm that this influx requires NRC4, we repeated the assay in *nrc2/3/4* knockout GCaMP6 plants. Wild-type NRC4 restored both the transient calcium influx and cell death, whereas the NRC4^L9E^ mutant did not (Fig. S5C), demonstrating that the observed responses are dependent on functional NRC4.

**Figure 3.**
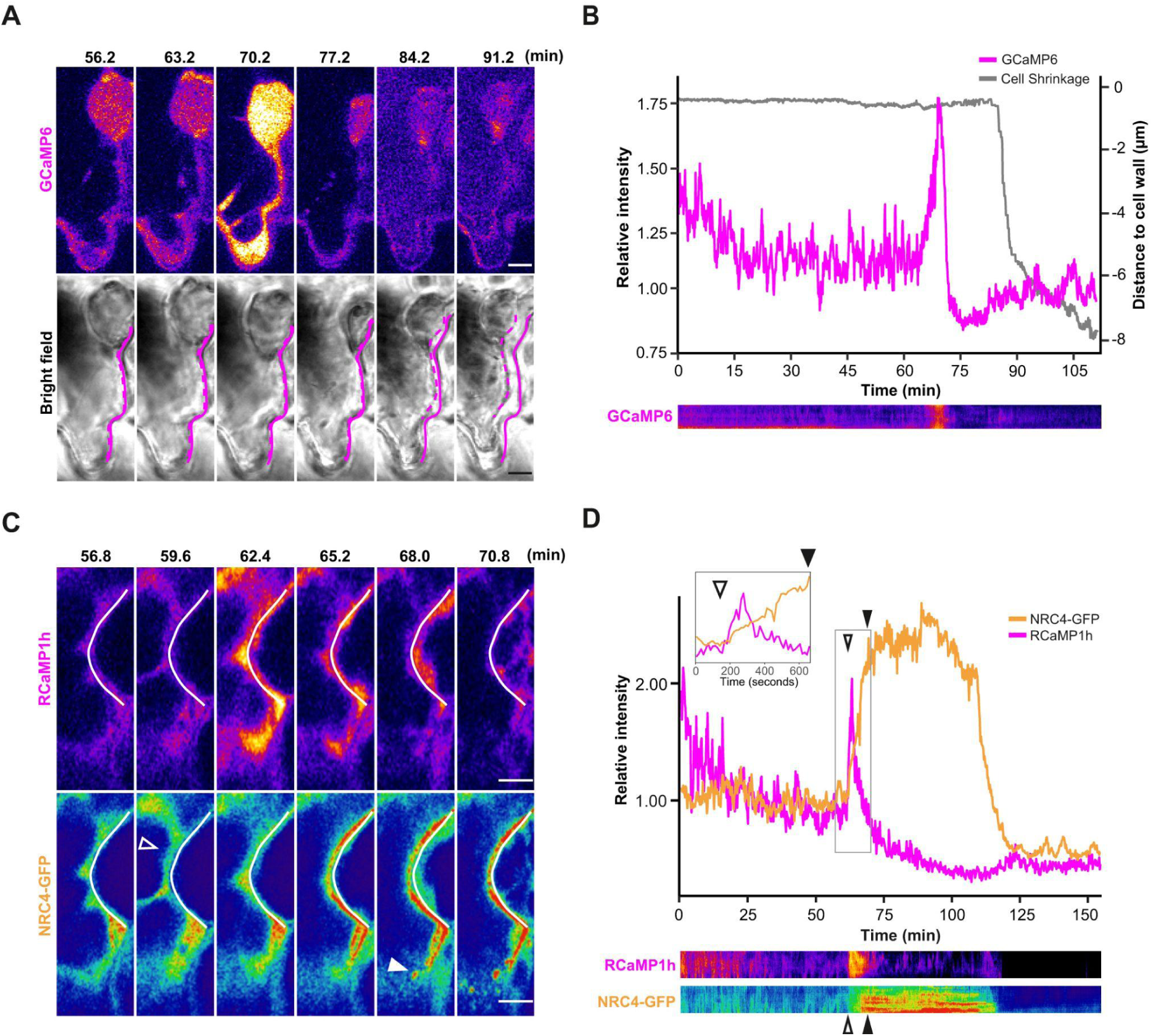
Activation of NRC4 resistosomes induces calcium influx into the cytoplasm. A. Time-lapse images showing cytosolic calcium dynamics during hypersensitive cell death. Leaves of GCaMP6-expressing *N. benthamiana* were infiltrated with *35S::Rpi-blb2*, *CBS4::AVRblb2*, and *35S::CUP2-p65*. Two days post infiltration, copper solution was applied to induce effector expression, followed by confocal imaging. Top: pseudocolored GCaMP6 fluorescence representing calcium intensity; bottom: corresponding bright-field images. Solid and dotted lines outline the right and the left cell boundaries. Images were extracted at 7-minute intervals from the time series in (B). Scale bar = 5 μm. (See Movie S5.) B. Quantification of GCaMP6 fluorescence intensity and plasma membrane shrinkage during hypersensitive cell death. GCaMP6 intensity was normalized to the mean intensity of 50 baseline frames recorded before calcium signal initiation. Membrane–cell wall distance was measured using the TrackMate plugin in ImageJ. Bottom: kymograph of the GCaMP6 signal from the time-lapse series in (A). C. Coordinated dynamics of cytosolic calcium (RCaMP1h) and plasma membrane-localized NRC4-GFP during hypersensitive cell death. *N. benthamiana* leaves were infiltrated with *35S::Rpi-blb2*, *CBS4::AVRblb2*, *35S::CUP2-p65*, *35S::RCaMP1h*, and *35S::NRC4-GFP*. White lines outline the cell boundaries. Top: pseudocolored RCaMP1h signal; bottom: pseudocolored NRC4-GFP signal. Open arrowheads mark initial NRC4 membrane enrichment; filled arrowheads indicate puncta formation. Images were extracted at 2.8-minute intervals from the time series. Scale bar = 5 μm. (See Movie S6.) D. Quantification of RCaMP1h and NRC4-GFP fluorescence intensities during cell death progression. Traces were normalized to the mean intensity of 50 baseline frames recorded before calcium signal initiation. The inset shows a zoomed-in time window around calcium signal onset and decline. Open and filled arrowheads denote NRC4 membrane enrichment and puncta formation, respectively. Bottom: kymographs of RCaMP1h and NRC4-GFP signals from the time-lapse series in (C).

To investigate the timing of calcium influx relative to NRC4 puncta formation, we co-imaged NRC4–GFP with the red-shifted calcium reporter RCaMP1h (Akerboom et al. 2013). To our surprise, visible NRC4 puncta appeared only after the calcium signal had already peaked (Fig. 3C, S4B; Movie S6). This prompted us to hypothesize that NRC4 resistosome are activated before puncta become detectable. To further investigate this, we quantified RCaMP and NRC4 fluorescence at the plasma membrane and found that the initial enrichment of NRC4 at the membrane coincided with the onset of calcium influx (Fig. 3D, S4B and Fig. S5D). NRC4 then reached a peak at the membrane and subsequently coalesced into puncta as the calcium signal declined. These data suggest that functional resistosomes assemble during the initial membrane-enrichment phase and that the later visible larger puncta represent post-activation clusters of multiple complexes rather than individual active resistosomes.

### NRC4 resistosome activation halts vesicle, mitochondrial, and Golgi dynamics

Because cytoplasmic streaming stops soon after NRC4 activation but before cell collapse, we asked whether other organelles show a comparable arrest. Since the formation of NRC4 resistosome cluster puncta is the most distinctive feature during the process, we used it as a reference point (time 0) to assess the timing of various subcellular events. We first examined ARA7-labeled endosomes and found that the movement of endosomes slowed markedly around 3 min before puncta became visible and had stopped entirely by that time, roughly matching the onset and peak of the transient calcium influx (Fig. 4A–B, and Fig. S6A; Movie S7) (Scheuring et al. 2011). Likewise, GmMan49-marked Golgi stacks lost motility ∼2–3 min before puncta formation (Fig. S7A,B; Movie S8) (Nelson et al. 2007).

**Figure 4.**
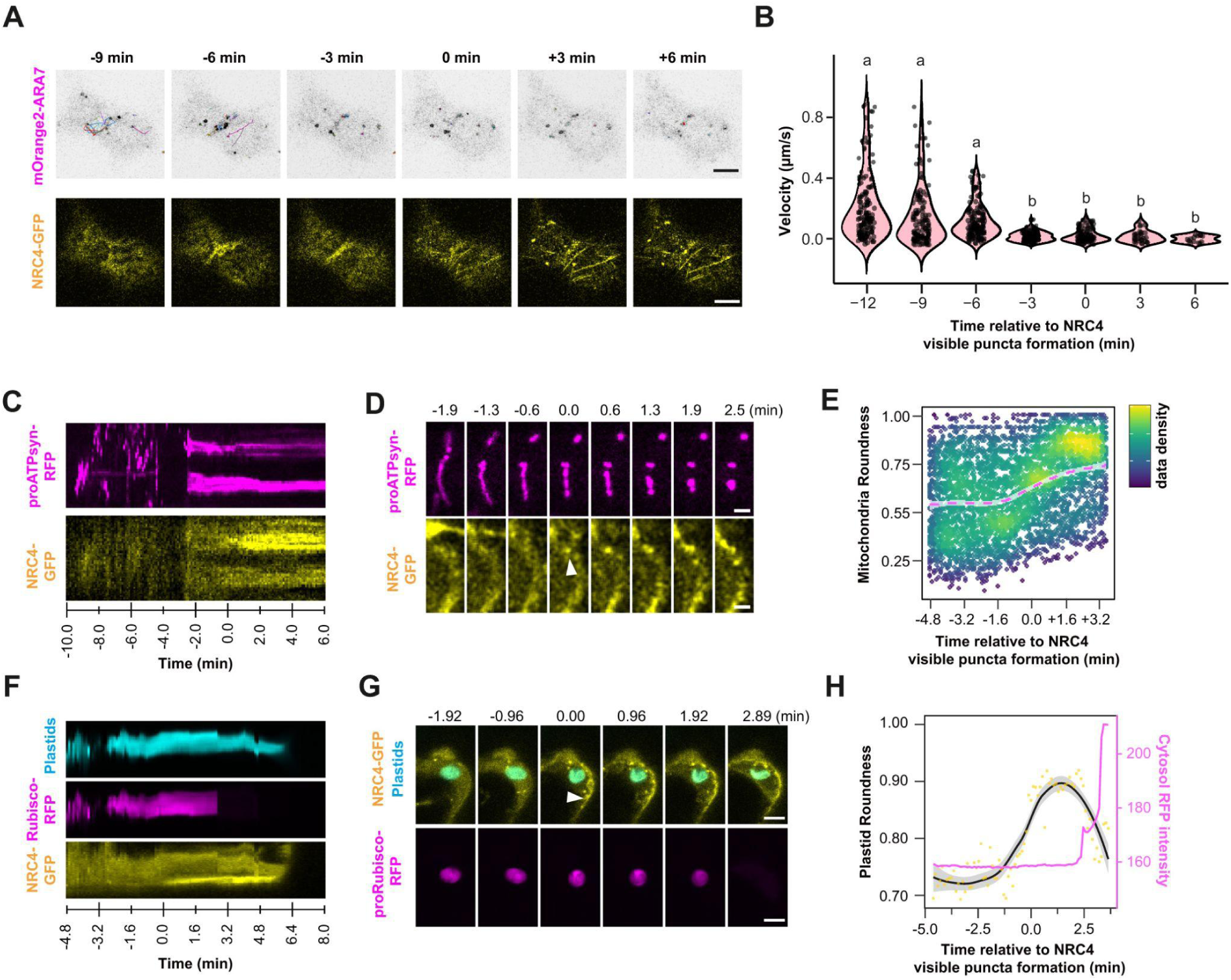
NRC4 resistosome activation halts the movement of vesicles, mitochondria, and Golgi bodies, and causes plastid disruption. A. Time-lapse confocal images showing the dynamics of mOrange2-ARA7–labeled endosomes (top) and NRC4-GFP (bottom) during hypersensitive cell death. *N. benthamiana* leaves were co-infiltrated with *35S::Rpi-blb2*, *CBS4::AVRblb2*, *35S::CUP2-p65*, *35S::mOrange2-ARA7*, and *35S::NRC4-GFP*. Two days post infiltration, copper was applied to induce effector expression. Images were acquired at 3-minute intervals. Time 0 corresponds to the onset of visible NRC4 puncta. Scale bar = 2.5 μm. (See Movie S7.) B. Quantification of endosome trafficking velocity at different time windows relative to NRC4 puncta formation. Velocities were calculated across 50-frame windows. Each dot represents an individual endosome; violin plots show distribution. Different letters denote statistically significant differences (one-way ANOVA with Tukey’s HSD, p < 0.05). C. Kymographs showing the temporal dynamics of mitochondria (proATPsyn-RFP, top) and NRC4-GFP (bottom) during hypersensitive cell death. D. Time-lapse images of mitochondrial morphology during NRC4 activation. Top: proATPsyn-RFP signal from the cell periphery; bottom: NRC4-GFP signal from the same region. Time 0 marks NRC4 puncta formation. White arrowhead indicates a representative NRC4 punctum. Scale bar = 2 μm. (See Movie S9.) E. Time-course quantification of mitochondrial roundness. Each point represents one mitochondrion. Color scale indicates data density (purple: low; yellow: high). The dashed line shows a generalized linear model (GLM) fit; shading represents the 95% confidence interval of the predicted mean. (n = 40-63 mitochondria.) F. Kymographs showing the temporal dynamics of plastids (autofluorescence, top), proRubisco-RFP–labeled stroma (middle), and NRC4-GFP (bottom) during hypersensitive cell death. G. Time-lapse images showing changes of plastid morphology and integrity during NRC4 activation. Top: merged image of plastid autofluorescence and NRC4-GFP; bottom: proRubisco-RFP signal. Time 0 marks visible NRC4 puncta formation. White arrowhead indicates an NRC4 punctum. Scale bar = 5 μm. (See Movie S10.) H. Quantification of plastid roundness and cytosolic proRubisco-RFP signal intensity over time. Only plastids that remained visible throughout the time series were analyzed. Yellow dots: individual plastid measurements; black line: GLM trend of predicted mean with 95% confidence interval (gray shading). Magenta line (right Y-axis): mean cytosolic proRubisco-RFP signal intensity. (n = 6-10 plastids.)

Time-lapse imaging using the mitochondrial matrix marker proATPsyn-RFP showed that mitochondria were highly motile and alternated between spherical and rod-like shapes (Lee et al. 2012). Roughly 2–3 min before NRC4 puncta appeared, mitochondrial movement stopped and the mitochondrial population became exclusively spherical, as reflected by a marked increase in roundness (Fig. 4C–E, and Fig. S6B; Movie S9). To examine plastid dynamics, we expressed the plastid stroma marker proRubisco-RFP and used chlorophyll autofluorescence to visualize thylakoids (Nelson et al. 2007). Similar to other organelles, plastids moved dynamically before NRC4 activation. Notably, around two minutes before the appearance of NRC4 puncta, plastids began to swell, followed by a rapid burst that released the stromal contents into the cytosol while the thylakoid signal remained intact (Fig. 4F-H, S6C; Movie S10). Overall, these observations indicate that NRC4 resistosome activation triggers dramatic changes in organelles, including cessation of movement, altered morphology, and loss of integrity. These events likely begin near the time of calcium influx, as inferred from Fig. 3, and continue during the appearance of NRC4 puncta, preceding cell collapse.

### NRC4 resistosome activation depolymerizes the actin cytoskeleton

We next tracked cytoskeletal dynamics during NRC4-mediated cell death. Live imaging with the LifeAct–mOrange2 marker revealed highly motile actin filaments under resting conditions. Their movement stopped around 3–4 min before NRC4 puncta became visible, and fluorescence then grew progressively weaker and more diffuse (Fig. 5A, S8A, and S9A–B; Movie S11). Quantitative anisotropy analysis of LifeAct-mOrange2-labeled actin filaments revealed a sharp drop in filament alignment at 0.5-1.0 minutes before the appearance of NRC4 puncta, indicative of rapid actin depolymerization (Fig. 5B). Similarly, we analyzed microtubule dynamics during NRC4-mediated cell death using the mOrange2-MAP4 marker. Anisotropy analysis revealed that microtubule filaments progressively lost alignment during cell death, with disassembly initiating 0.5-1.0 minutes before the appearance of NRC4 puncta (Fig. 5C-D, S8B, and S9C–D; Movie S12). Intriguingly, NRC4 filaments arose in regions that previously had been occupied by microtubules, although any functional relationship between microtubules and NRC4 resistosomes remains so far unclear (Movie S13). Because the cytoskeleton underpins the motility and positioning of multiple organelles, its disassembly could explain the abrupt arrest of organelle movement and may help remodel cellular architecture for the rapid execution of the immune response (Perico and Sparkes 2018).

**Figure 5.**
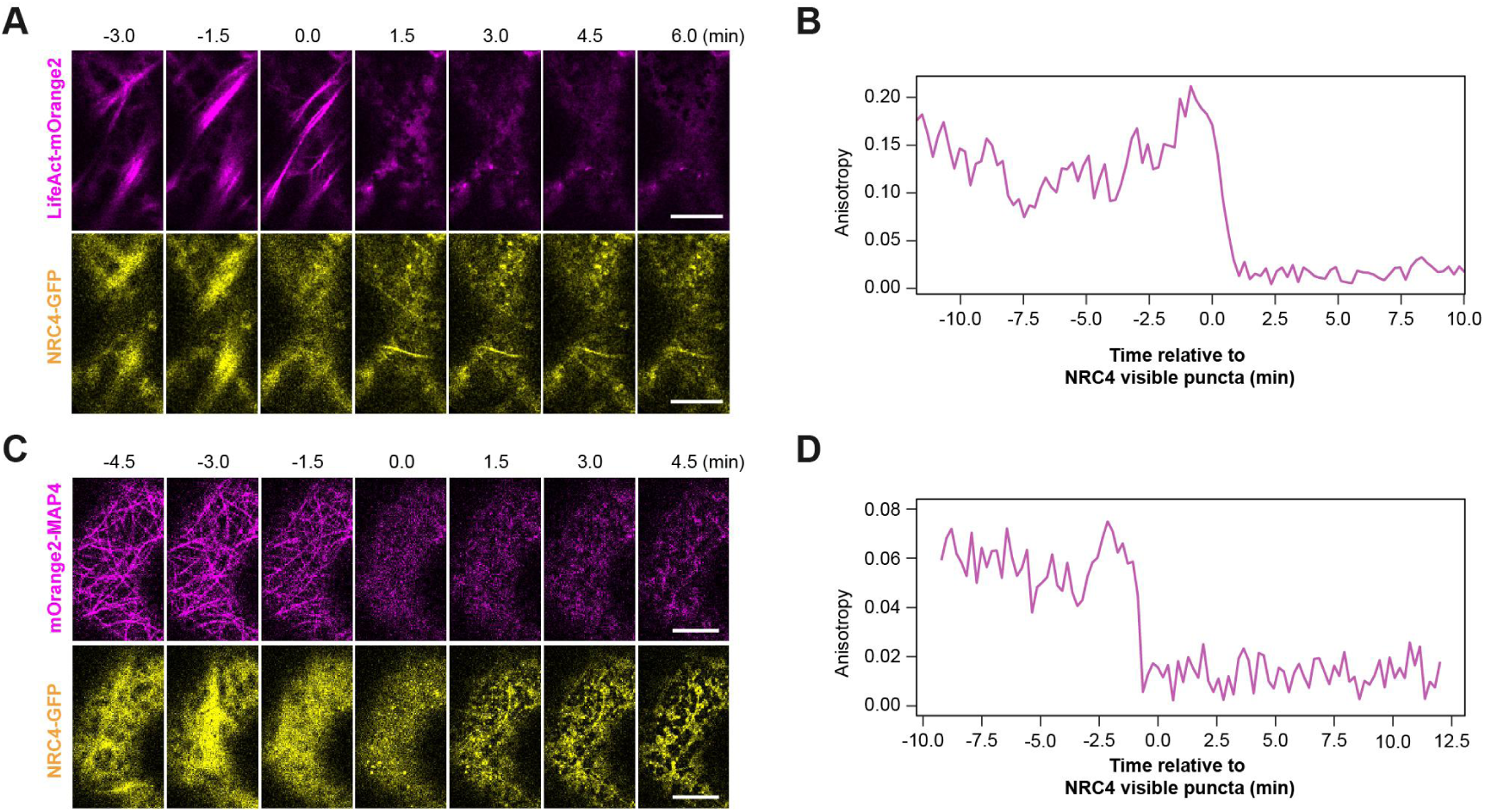
Cytoskeletons undergo depolymerization upon NRC4 resistosome activation. A. Time-lapse images showing actin dynamics labeled by LifeAct-mOrange2 (top) and NRC4-GFP (bottom) during hypersensitive cell death. *N. benthamiana* leaves were infiltrated with *35S::Rpi-blb2*, *CBS4::AVRblb2*, *35S::CUP2-p65*, *35S::LifeAct-mOrange2*, and *35S::NRC4-GFP*. Copper solution was applied 2 days post infiltration, followed by confocal imaging. Images were subsetted from the time series at 1.5-minute intervals. Time 0 corresponds to the first appearance of NRC4 puncta. Scale bar = 10 μm. (See Movie S11.) B. Quantification of actin filament anisotropy over time. Filament coverage was measured within the cortical region and expressed as anisotropy values indicating directional alignment. A rapid decline in actin anisotropy was observed upon NRC4 puncta formation. C. Time-lapse images showing microtubule dynamics labeled by mOrange2-MAP4-MBD (top) and NRC4-GFP (bottom) during hypersensitive cell death. Plants were co-infiltrated with *35S::Rpi-blb2*, *CBS4::AVRblb2*, *35S::CUP2-p65*, *35S::mOrange2-MAP4-MBD*, and *35S::NRC4-GFP*. Copper was applied at 2 dpi, and images were subsetted from the time series at 1.5-minute intervals. Time 0 corresponds to visible NRC4 puncta formation. Scale bar = 10 μm. (See Movie S12.) D. Quantification of microtubule anisotropy over time. Filamentous regions were segmented within the cortical plane, and anisotropy was computed as a measure of microtubule organization. Microtubule alignment declined sharply following NRC4 puncta formation.

### Activation of NRC4 resistosome leads to loss of ER, plasma membrane, and tonoplast integrity

Since an intact cytoskeleton helps maintain endoplasmic-reticulum (ER) morphology, we tracked ER dynamics during NRC4-mediated cell death using the ER marker mCherry-HDEL (Wang and Hussey 2015; Pain et al. 2023). Initially, the ER exhibited dynamic movement during the early phases of imaging after copper treatment. About four minutes before NRC4 puncta appeared, this movement stopped, although the reticulate network remained intact. Strikingly, once NRC4 puncta became visible, the ER network fragmented into vesicle-like structures (Fig. 6A and S10A; Movie S14). Persistence mapping confirmed that the network was stable during the ten minutes preceding puncta formation (Fig. S11A; Movie S14), but disintegrated during the following ten minutes(Fig. S11AB; Movie S14). Quantitative analysis showed that ER mesh counts fell from around 15-20 per 400 μm^2^ to zero at the moment NRC4 puncta appeared, confirming the swift transition from a tubular network to dispersed vesicles-like forms (Fig. 6B).

**Figure 6.**
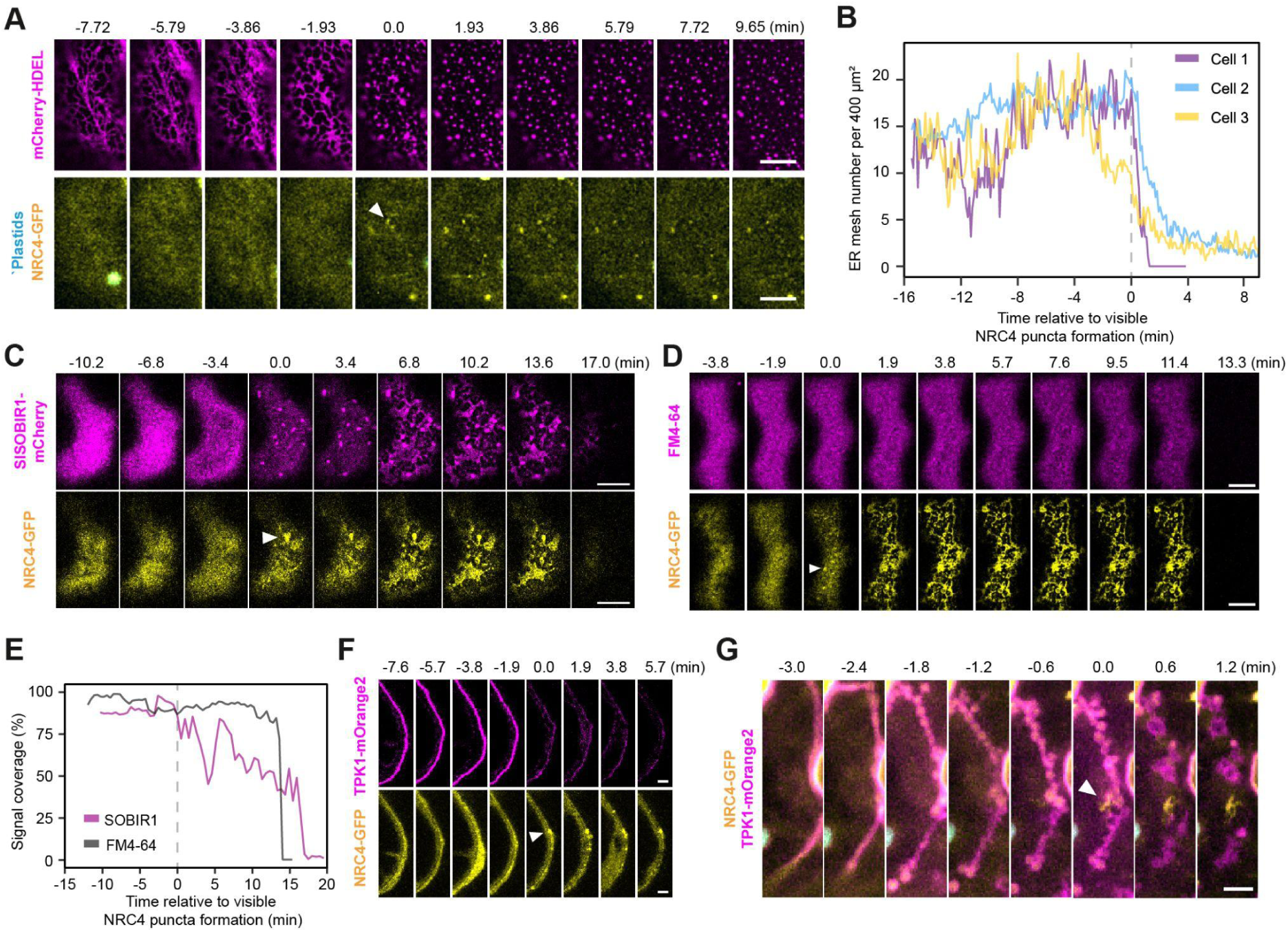
NRC4 resistosome activation disrupts ER, plasma membrane, and tonoplast integrity. A. Time-lapse images showing fragmentation of the endoplasmic reticulum (ER) during hypersensitive cell death. *N. benthamiana* leaves were co-infiltrated with *35S::Rpi-blb2*, *CBS4::AVRblb2*, *35S::CUP2-p65*,*35S::mCherry-HDEL, and 35S::NRC4-GFP*. Two days post infiltration, copper solution was applied, followed by confocal imaging. Top: mCherry-HDEL; bottom: NRC4-GFP. Images were extracted from the time series at 4.2-minute intervals. White arrowhead marks an NRC4 punctum at the cell periphery. Scale bar = 10 μm. (See Movie S14.) B. Quantification of ER mesh structures over time. Meshes were manually segmented in cortical regions from three cells, where a mesh was defined as an area enclosed by ER tubules. Time 0 corresponds to the appearance of NRC4 puncta at the cell periphery. C. and D. Time-lapse images showing changes in PM protein and lipid distribution during hypersensitive cell death. Leaves were co-infiltrated as in (A), and additionally with *SlSOBIR1-mCherry* for PM protein labeling (C), or stained with FM4-64 for lipid labeling (D). *NRC4-GFP* was co-expressed in both setups. Images were extracted from the time series at 3.4-minute (C) and 1.9-minute (D) intervals. White arrowheads indicate NRC4 puncta formation. Scale bars = 10 μm. (See Movies S15, S16.) E. Quantification of plasma membrane coverage by SlSOBIR1-mCherry and FM4-64 over time. Signal coverage is expressed as the percentage of membrane area labeled relative to the initial frame. Time 0 marks the onset of visible NRC4 puncta. F. Time-lapse images showing tonoplast dynamics using AtTPK1-mOrange2 as a tonoplast marker. Plants were co-infiltrated with *35S::Rpi-blb2*, *CBS4::AVRblb2*, *35S::CUP2-p65*, *AtTPK1-mOrange2, and 35S::NRC4-GFP*. Representative images were extracted at a 1.9-minute interval. Top: TPK1-mOrange2 signal; bottom: NRC4-GFP signal. White arrowhead indicates an NRC4 punctum. Scale bar = 5 μm. (See Movie S17.) G. Maximum intensity projection showing merged signals of NRC4-GFP and AtTPK1-mOrange2 during disruption of trans-vacuolar strands. Time 0 marks the point of visible NRC4 puncta formation. White arrowhead indicates an NRC4 punctum at the cell edge. Scale bar = 5 μm. (See Movie S19.)

To visualise plasma-membrane behaviour during NRC4-mediated cell death, we used SlSOBIR1-mCherry and the lipophilic dye FM4-64 (Peng et al. 2015). As soon as NRC4-GFP puncta became visible, SlSOBIR1-mCherry reorganised into discrete, protein-rich microdomains; around 6–7 min later these microdomains resolved into a reticulate pattern, implying a profound change in membrane organization (Fig. 6C; Movie S15). In contrast, the bulk lipid distribution labelled by FM4-64 showed little change over the same period (Fig. 6D; Movie S16). Quantification confirmed that SlSOBIR1 coverage at the plasma membrane declined steadily after NRC4 activation, whereas FM4-64 coverage remained stable until the late phase when overt cell collapse became apparent (Fig. 6E; Movie S16). These findings indicate that NRC4 resistosome activation alters the protein landscape and biophysical properties of the plasma membrane well before the lipid bilayer itself is compromised.

To investigate tonoplast dynamics, we used AtTPK1 as a marker for the tonoplast membrane (Kasaras and Kunze 2017). The AtTPK1-mOrange2 signal remained uniform until NRC4 puncta appeared, after which fluorescence intensity and membrane coverage dropped sharply, signalling rapid loss of tonoplast integrity (Fig. 6F, S10B–C, and S11D; Movies S17, S18). Roughly two minutes before puncta formation, transvacuolar strands, whose stability typically depends on the cytoskeleton, began to bulge into bleb-like swellings; these strands then fragmented into vesicle-like bodies and gradually vanished (Fig. 6G; Movie S19). These observations suggest that NRC4-mediated hypersensitive cell death involves coordinated remodeling and destabilization of multiple membrane systems, including the tonoplast.

### NRC4 resistosome activation triggers nuclear shrinkage and nucleoplasm release before cell collapse

Nuclear shrinkage is a hallmark of cell death (Mur et al. 2008). To track the dynamics of the nucleus during NRC4-mediated cell death, we expressed NRC4-GFP along with BFP-NLS and performed PI staining. Time-lapse experiments revealed that nuclear shrinkage started at the same time or shortly before the appearance of NRC4 puncta (Fig. 7A-B, S7A-B; Movie S20, S21). As contraction continued, BFP-NLS leaked into the cytosol, indicating loss of nuclear-envelope integrity (Fig. 7A; Movie S20). Initially, the released BFP signal was restricted to the cytosol, indicating that the tonoplast was still intact at that stage. The BFP signal then abruptly dispersed into the center of the cell, likely reflecting the rupture of the tonoplast membrane, and then quickly diminished thereafter.(Fig. 7A-B, S12, and S13A–C; Movie S20, S21). PI entered the nucleus at the moment NRC4 puncta formed and steadily increased in intensity, saturating ∼20–25 min later (Fig. 7A-B, S13C; Movie S20, S21). Together, these data place nuclear collapse and tonoplast rupture downstream of NRC4 activation but upstream of the final cellular collapse.

**Figure 7.**
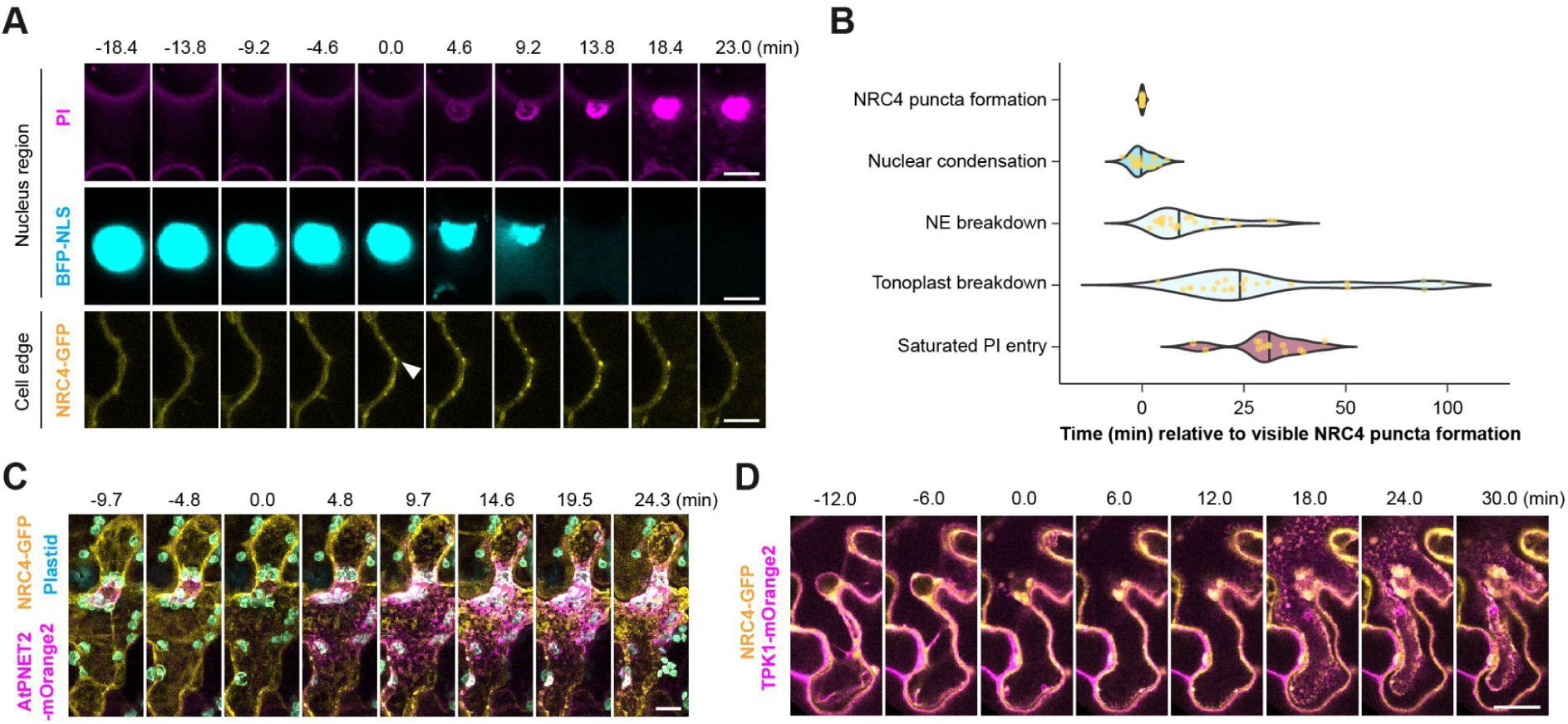
NRC4 resistosome activation induces nuclear condensation and nucleoplasm release, followed by cell collapse. A. Dynamic changes in nuclear integrity, morphology and NRC4 localization during hypersensitive cell death. *N. benthamiana* leaves were agro-infiltrated with *35S::Rpi-blb2*, *CBS4::AVRblb2*, *35S::CUP2-p65*, *35S::BFP-NLS* and *35S::NRC4-GFP*. At 2 days post-agroinfiltration, copper solution and propidium iodide were applied, followed by confocal imaging. The montage was generated from time-lapse images with 4.6-minute intervals. Top and middle panels show the signals of PI staining and BFP-NLS fluorescence, respectively; the bottom panel shows NRC4-GFP fluorescence signal cropped from the cell edge. Time zero was set to be relative to NRC4 puncta formation, with white arrowheads indicating a visible NRC4 punctum. Scale bar = 10 μm. (Movie S20) B. Temporal analysis of subcellular events occurring between NRC4 puncta formation and saturated PI entry. Each dot represents the time point for a subcellular event occurring in a cell. Time points for nuclear condensation, nuclear envelope (NE) breakdown, and tonoplast breakdown, were determined from the time-lapse series of n = 23 cells. For PI entry, n = 16 cells were analyzed, with 3 outliers excluded from the plot. The x-axis shows the time relative to NRC4 puncta formation (set as time zero). C. Time-lapse montage showing the dynamics of the nuclear envelope label with AtPNET2 during cell death. *N. benthamiana* leaves were agro-infiltrated with *35S::Rpi-blb2*, *CBS4::AVRblb2*, *35S::CUP2-p65*, *35S::AtPNET2-mOrange2* and *35S::NRC4-GFP*. At 2 days post-agroinfiltration, copper solution was applied, followed by confocal imaging. The montage was generated from time-lapse 3D image stacks captured at 4.8-minute intervals, showing merged signals from three fluorescence channels. Scale bar = 10 μm. (Movie S22) D. Time-lapse montage showing tonoplast dynamics during hypersensitive cell death. Leaves were agro-infiltrated with *35S::Rpi-blb2*, *CBS4::AVRblb2*, *35S::CUP2-p65*, *35S::TPK1-mOrange2* and *35S::NRC4-GFP*. At 2 days post-agroinfiltration, copper solution was applied, followed by confocal imaging. The montage was generated from time-lapse images with 6-minute intervals. Scale bar = 20 μm. (Movie S23)

Because nucleoplasm leakage implied loss of nuclear envelope integrity, we monitored the envelope directly with the outer-nuclear-membrane marker AtPNET2-mOrange2 (Tang et al. 2022). Prior to the appearance of NRC4 puncta, AtPNET2-mOrange2 sharply outlined the nuclear rim (Fig. 7C). Approximately 4–5 minutes after NRC4 puncta formation, the AtPNET2-mOrange2 signal began to diffuse into the cytosol, indicating nuclear envelope breakdown (Fig. 7C; Movie S22). In the final collapse phase—typically 20–30 min after puncta emergence, though the timing varied across individual cells—the tonoplast and plasma membrane, detached from the cell wall and retracted toward the cell center (Fig. 7D, and Fig.S14; Movie S23).

## Discussion

Using a copper-inducible expression system (Chiang et al. 2024), we conducted high-resolution time-lapse imaging to capture the sequence of subcellular events that underlie resistosome-mediated hypersensitive cell death, a process that has been very challenging to visualize in the past (Fig. S15). Our analyses revealed that NRC4 resistosome activation triggers a rapid influx of calcium into the cell, coinciding with NRC4 enrichment at the plasma membrane. This spike initiates a cascade of profound cellular changes: organelle motility ceases, morphology is disrupted, and membrane integrity deteriorates. Both actin filaments and microtubules depolymerise rapidly, and membrane systems remodel on a broad scale: plasma-membrane proteins redistribute, the ER and transvacuolar structures fragment into vesicle-like bodies, and the tonoplast loses integrity. Nuclear events begin with shrinkage, followed by nucleoplasm leakage and rupture of the nuclear envelope. These processes culminate in catastrophic tonoplast failure and complete cell collapse, marked by the plasma membrane peeling away from the cell wall.

Since calcium influx can trigger actin and microtubule depolymerization (Cai et al. 2015; Madina et al. 2019), we propose that cytoskeleton breakdown is a coordinated component of the death execution program, facilitating the architectural dismantling associated with immune-mediated cell collapse. However, additional, as yet unidentified, signaling cascades are likely to contribute to the physiological transitions and final stages of cell death. Together, our findings provide a spatiotemporally resolved framework for understanding the comprehensive subcellular reorganization driven by resistosome activation, shedding light on how immune receptors orchestrate hypersensitive cell death.

A key next step is to determine how the calcium influx triggered by resistosome activation is decoded within the cell. It remains unclear whether calcium alone is sufficient to orchestrate the diverse subcellular processes observed during resistosome-mediated cell death, or whether additional secondary messengers are required. Furthermore, it is not known whether certain organelles play more central roles in mediating cell death, or whether the observed changes are merely downstream consequences of the death process. Addressing these questions will be crucial for understanding how immune signals are integrated at the cellular level to coordinate cell death and defense.

Recent transcriptomic studies have begun to identify genetic components associated with immune-triggered cell death (Salguero-Linares et al. 2022; Burke et al. 2023). Although based on different experimental systems, it would be valuable to investigate whether these candidate genes influence the organelle dynamics described here. Additionally, enzymes such as metacaspases and autophagy-related proteins, both previously implicated in plant hypersensitive cell death (Hofius et al. 2009, 2017; Coll et al. 2014), may act at specific stages such as cytoskeletal disassembly or membrane rupture, or may operate more broadly across death cascade.

This study focuses on the NRC4 resistosome, a hexameric complex representing one of several structural classes of resistosomes in plants (Huang et al. 2025a). Other forms, such as pentameric resistosomes, likely function in a similar manner, while TNL-derived tetrameric resistosomes act as NADase enzymes to induce cell death through distinct mechanisms (Huang et al. 2025a). Whether these structurally diverse complexes, as well as downstream partners such as NRG1 and ADR1, converge on common or distinct cell death programs remains an open question (Yu et al. 2024; Huang et al. 2025b). Similarly, the differences between the cell death processes observed here to the vacuolar-type cell death mediated by proteins like PML5, or membrane-disrupting cell death triggered by MLKL-like proteins, require further investigation (Shen et al. 2024; Sunil et al. 2024). Beyond NLRs, cell surface immune receptors such as Cf-4 and ELR also induce cell death upon recognition of the corresponding ligands (Thomas et al. 1997; Du et al. 2015). Although NRCs, in particular NRC3, have been shown to contribute to cell death triggered by cell-surface immune receptors, the extent of overlap in downstream signaling, including calcium dynamics and organelle remodeling, between PRR- and NLR-mediated pathways remains unclear (Kourelis et al. 2022).

Finally, programmed cell death during plant development shares morphological features with immune-related cell death, but the mechanistic parallels remain underexplored (Wang et al. 2023). Beyond the plant kingdom, animal cell death programs such as apoptosis and pyroptosis exhibit distinct but occasionally analogous features (Coll et al. 2011; Maekawa et al. 2023). How much convergence exists in subcellular dynamics between plant and animal immune cell death programs remains to be addressed (Mur et al. 2008). Our spatiotemporally resolved analyses of calcium signaling and organelle dynamics offer a valuable reference framework for further investigations into the mechanisms of plant cell death across immune, evolutionary, and developmental contexts, and also inform broader understanding of cell death processes beyond the plant kingdom.

## Acknowledgments

We thank Mark Youles (SynBio, The Sainsbury Laboratory, UK) for sharing plasmids for molecular cloning, Dr. Tien-Shin Yu (Institute of Plant and Microbial Biology, Academia Sinica), and Dr. Yen-Ping Hsueh (Institute of Molecular Biology, Academia Sinica) for sharing markers for cell biology studies. We thank Mei-Jane Fang and Ji-Ying Huang in the Cell Biology Core Lab (Institute of Plant and Microbial Biology, Academia Sinica, Taiwan) and Shu-Chen Shen in the Advanced Optical Microscope Core Facility (Agricultural Biotechnology Research Center, Academia Sinica, Taiwan) for help with confocal imaging. We thank Lin-Yun Kuang in the Transgenic Plant Laboratory (Academia Sinica) for generating transgenic *N. benthamiana* lines.

## Author contributions

Conceptualization: YFC, KYL, CYH, CHW

Methodology: YFC, KYL, CYH, HYW

Investigation: YFC, KYL, CYH, LYH, WCS, BJC, CWC

Visualization: YFC, KYL

Funding acquisition: CHW, TOB

Project administration: CHW

Supervision: CHW, TOB

Writing – original draft: YFC, KYL, LYH, BJC, ELHY, CHW

Writing – review & editing: YFC, KYL, LYH, CYH, WCS, BJC, ELHY, TOB, CHW

## Data and materials availability

All data are available in the main text or the supplementary materials.

---

**Figure S1.**
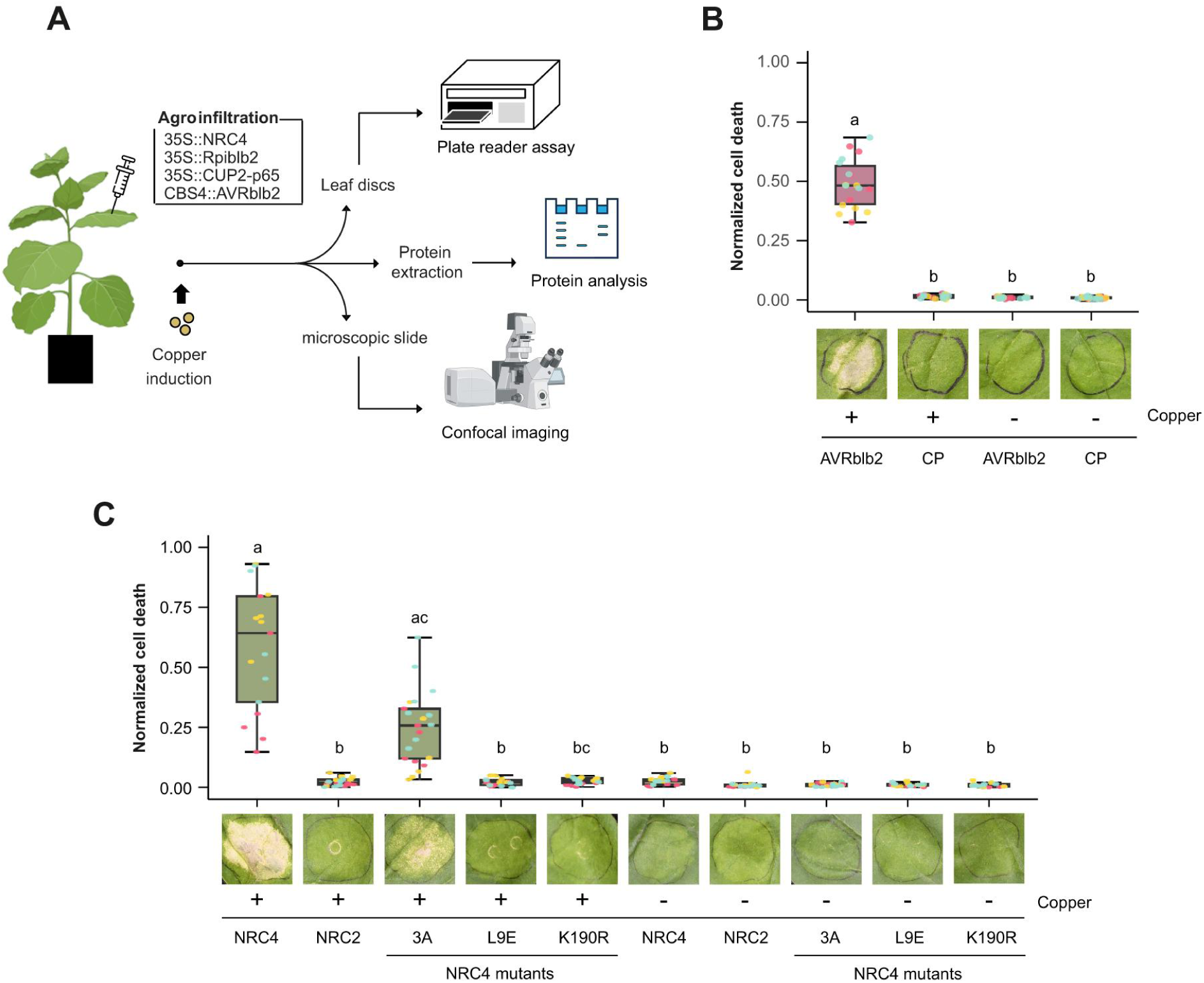
Effector expression driven by the copper inducible system triggers hypersensitive cell death in *N. benthamiana*. A. Schematic representation of the experimental workflow. *NRC4*, *Rpi-blb2*, and *CUP2-p65* were transiently expressed in *N. benthamiana* leaves under the control of *35S* promoter together with a copper-inducible *AVRblb2*. At two days post agro-infiltration, copper solution was infiltrated into leaves followed by plate reader assay, protein analysis and confocal imaging. B. The cell death phenotype of leaves expressing AVRblb2 or CP in the presence or absence of copper. Leaves of *N. benthamiana* were agro-infiltrated with *35S::Rpi-blb2, 35S::CUP2-p65,* and either *CBS4::AVRblb2* or *CP*. At 24 hours post-agroinfiltration, the copper solution was infiltrated into leaves. Autofluorescence emitted from dead cells were recorded using UVP ChemStudio at 36 h after copper infiltration. Raw fluorescence intensity was normalized to the maximum detectable value to represent normalized cell death levels. Dots with different colors represent the results from three independent biological replicates (n = 18). C. The cell death phenotype of leaves expressing NRC2, NRC4 or NRC4 mutants in the presence or absence of copper. Leaves of *N. benthamiana nrc* plants were agro-infiltrated with *35S::Rpi-blb2, CBS4::AVRblb2*, *35S::CUP2-p65,* and either *35S::NRC2*, *35S::NRC4* or other NRC4 mutants. At 24 hours post-agroinfiltration, the copper solution was infiltrated into leaves Autofluorescence emitted from dead cells were recorded using UVP ChemStudio at 36 h after copper infiltration. Raw fluorescence intensity was normalized to the maximum detectable value to represent normalized cell death levels. Raw fluorescence intensity was normalized to the maximum detectable value. Dots with different colors represent the results from three independent biological replicates (n = 18).

**Figure S2.**
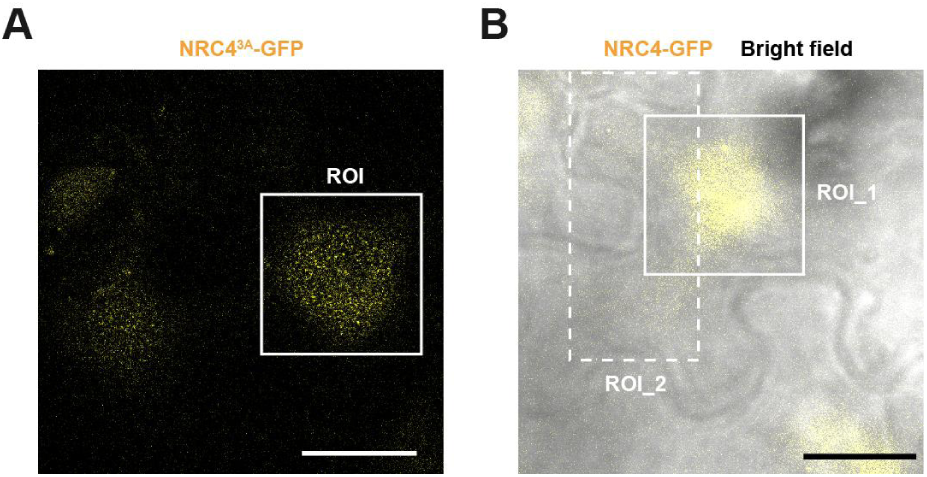
Regions of interest selected for generating montages in Figure 2. A. Representative NRC4^3A^-GFP fluorescence image. The white solid box indicates the region of interest (ROI) used to generate the time-lapse montages shown in Fig. 2A. Scale bar = 20 μm. B. Overlay of NRC4-GFP fluorescence and bright-field image. The solid and dashed boxes denote the regions corresponding to ROI_1 and ROI_2, which were used for time-lapse analysis presented in Fig. 2B and 2E, respectively. Scale bar = 20 μm.

**Figure S3.**
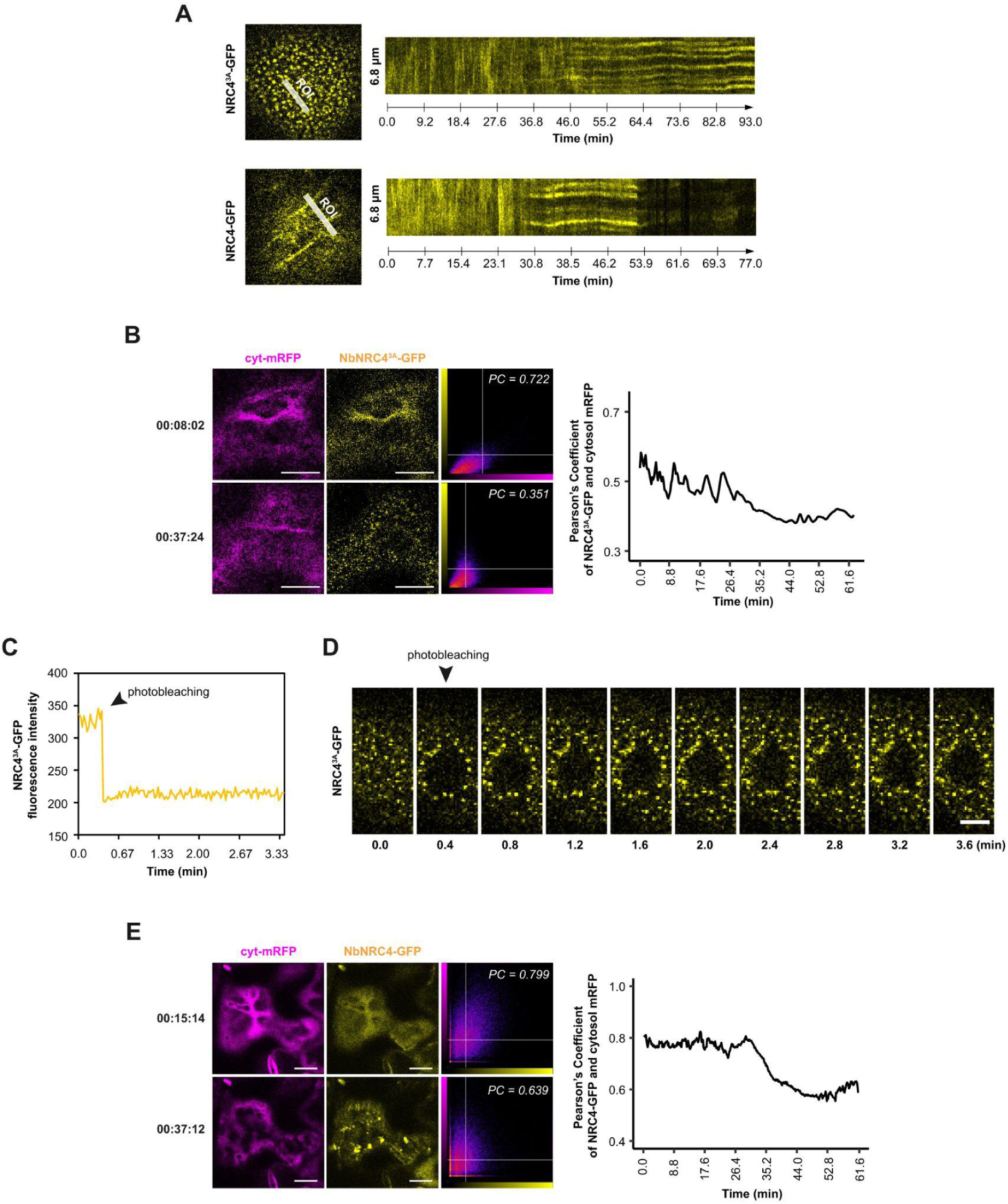
NRC4 spatially separated from cytosol upon activation. A. Kymographs showing the temporal dynamics of NRC4^3A^-GFP (top) and NRC4-GFP (bottom) in *N. benthamiana nrc* plants after copper-induced resistosome activation. Leaves were co-infiltrated with *35S::Rpi-blb2*, *CBS4::AVRblb2*, *35S::CUP2-p65*, and either *NRC4^3A^-GFP* or *NRC4-GFP*. Confocal imaging was performed 2 days post infiltration. White lines indicate regions of interest (ROIs) used for kymograph analysis, displayed on the right. B. Colocalization analysis of NRC4^3A^-GFP and cytosolic mRFP. Left: representative images of cells before (top) and after (bottom) NRC4^3A^-GFP puncta formation. Insets show scatterplots of pixel intensity correlations (NRC4^3A^-GFP: yellow, Y-axis; mRFP: magenta, X-axis) with corresponding Pearson’s correlation coefficients. Right: time-course of Pearson’s correlation coefficients between NRC4^3A^-GFP and cyt-mRFP signals. Scale bar = 10 μm. (See Movie S2.) C. Fluorescence recovery after photobleaching (FRAP) analysis of NRC4^3A^-GFP puncta. The arrowhead marks the photobleaching event. Fluorescence intensity was measured over time at 1.6-second intervals. D. Time-lapse images from the FRAP assay shown in (C), displayed at 24-second intervals. The arrowhead indicates the photobleaching timepoint; the white circle denotes the bleached area. Scale bar = 5 μm. E. Colocalization analysis of NRC4-GFP and cytosolic mRFP. Left: representative images before (top) and after (bottom) NRC4 activation. Scatterplots show intensity correlation between NRC4-GFP (yellow) and cyt-mRFP (magenta), with Pearson’s coefficients indicated. Right: time-course of Pearson’s correlation coefficients during NRC4 activation. Scale bar = 10 μm. (See Movie S4.)

**Figure S4.**
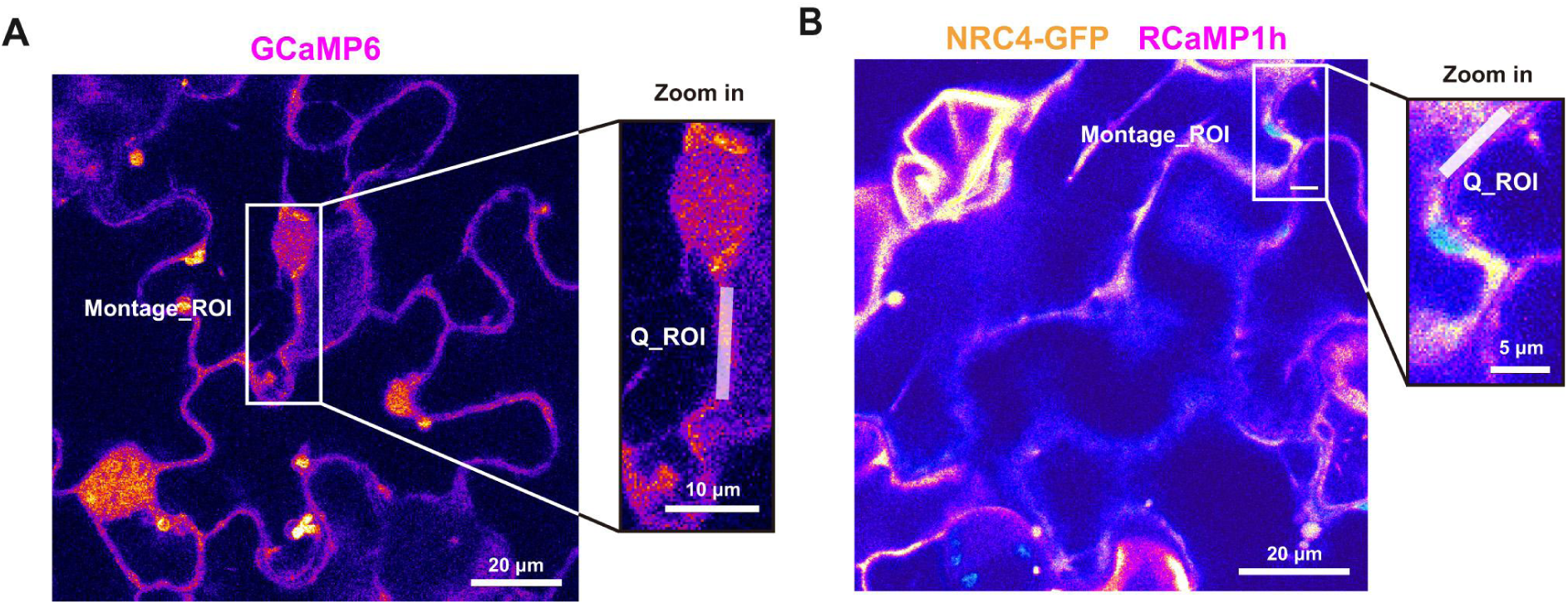
Regions of interest selected for generating montages and quantifying intensity in Figure 3. A. and B. Representative confocal images of GCaMP6 (A) and overlaid RCaMP1h/NRC4-GFP (B), corresponding to Movies S5 and S6, respectively. Left panels: original fields of view with white boxes indicating the cell regions used for montages in Fig. 3A. and 3C. Right panels: zoomed-in views with white lines denoting the regions of interest (ROIs) used for quantifying (Q_ROI) fluorescence intensity and generating kymographs shown in Fig. 3B. and 3D.

**Figure S5.**
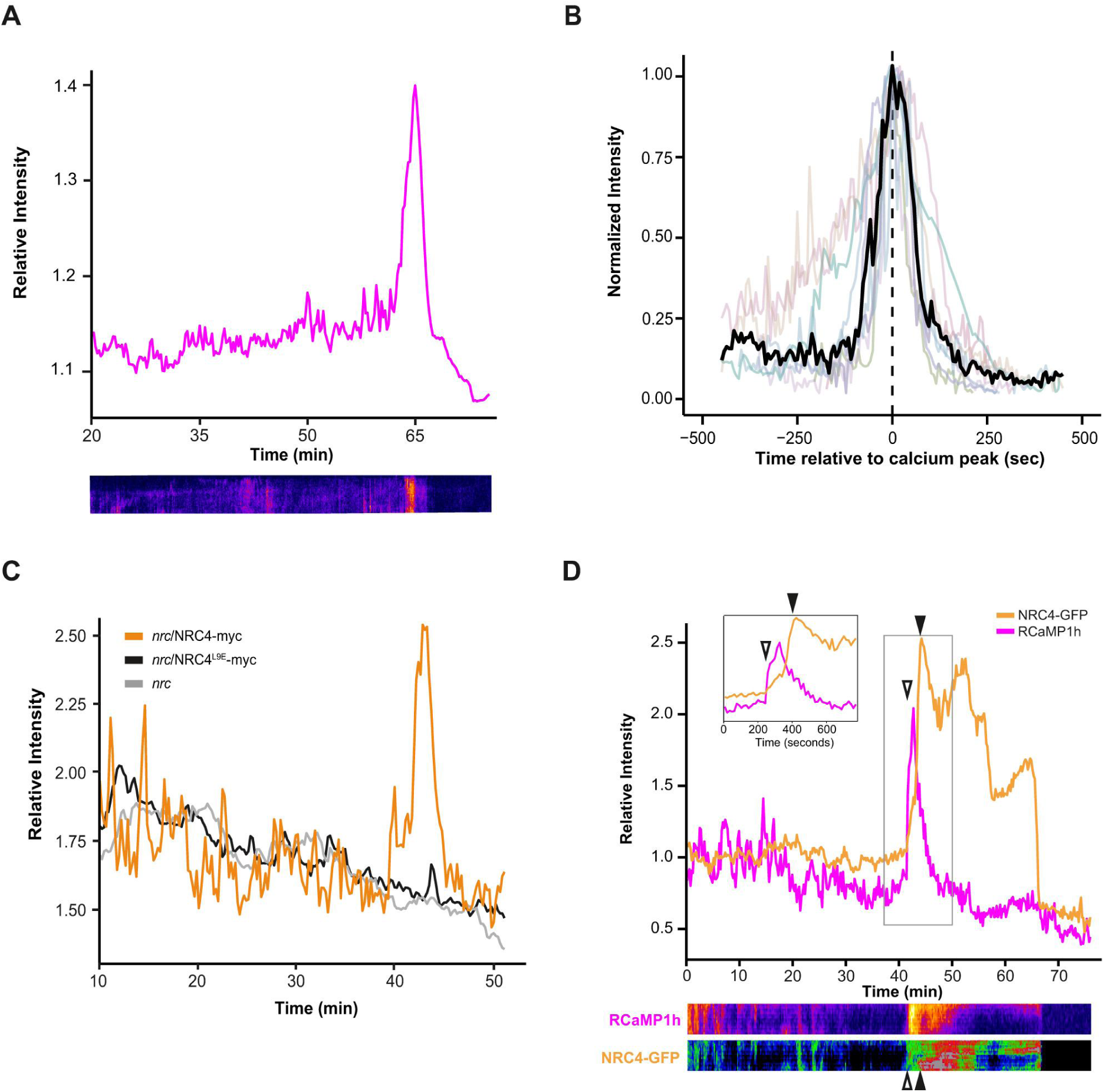
Co-expression of Rpi-blb2 and AVRblb2 triggers NRC4-dependent calcium influx. A. Quantification of cytosolic GCaMP6 intensity from a cell independent of cell shown in Fig. 3A. Leaves of GCaMP6-expressing *N. benthamiana* were infiltrated with *35S::Rpi-blb2*, *CBS4::AVRblb2*, *35S::CUP2-p65*. Two days post-agroinfiltration, copper solution was applied to induce effector expression, followed by confocal imaging. Time-lapse images were acquired at 6.43-second intervals, starting at 20 min after copper treatment. GCaMP6 intensity was normalized to the mean intensity of 50 baseline frames recorded before calcium signal initiation. Bottom: kymograph of the GCaMP6 signal from the time-lapse series. B. Alignment of calcium influx from independent cells. Calcium levels, indicated by GCaMP6 intensity, were normalized to the peak intensity of each time series. Data from independent cells were aligned to time 0, corresponding to the peak intensity, as a reference point. Lines represent individual cells. The black line represents the cell with the median progression timing of all cells analyzed (n = 10 cells). C. Quantification of GCaMP6 fluorescence intensity in *nrc2/3/4* knockout GCaMP6 plants in the presence or absence of NRC4 complementation. Leaves of *N. benthamiana* were infiltrated with *35S::Rpi-blb2*, *CBS4::AVRblb2*, *35S::CUP2-p65* and either *35S::NRC4*, *35S::NRC4*^L9E^ or without complementation. Two days post-agroinfiltration, the copper solution was infiltrated into leaves followed by confocal microscopy analysis. Time-lapse images were acquired at 6.43-second intervals after copper treatment. D. Quantification of RCaMP1h and NRC4-GFP fluorescence intensity during hypersensitive cell death from a cell independent of cell shown in Fig. 3C. Leaves of *N. benthamiana* were infiltrated with *35S::Rpi-blb2*, *CBS4::AVRblb2*, *35S::CUP2-p65*, *35S::RCaMP1h* and *35S::NRC4-GFP*. At 2 days post-agroinfiltration, the copper solution was applied to leaves followed by confocal microscopy analysis. Time-lapse images were acquired at 12.86-second intervals. RCaMP1h intensity was normalized to the mean intensity of 50 baseline frames recorded before calcium signal initiation. The inset at the top left shows a shorter time window spanning the onset and decline of calcium signals. Open arrowheads mark initial NRC4 membrane enrichment; filled arrowheads indicate puncta formation. Bottom: kymographs of RCaMP1h and NRC4-GFP signals from the time-lapse series.

**Figure S6.**
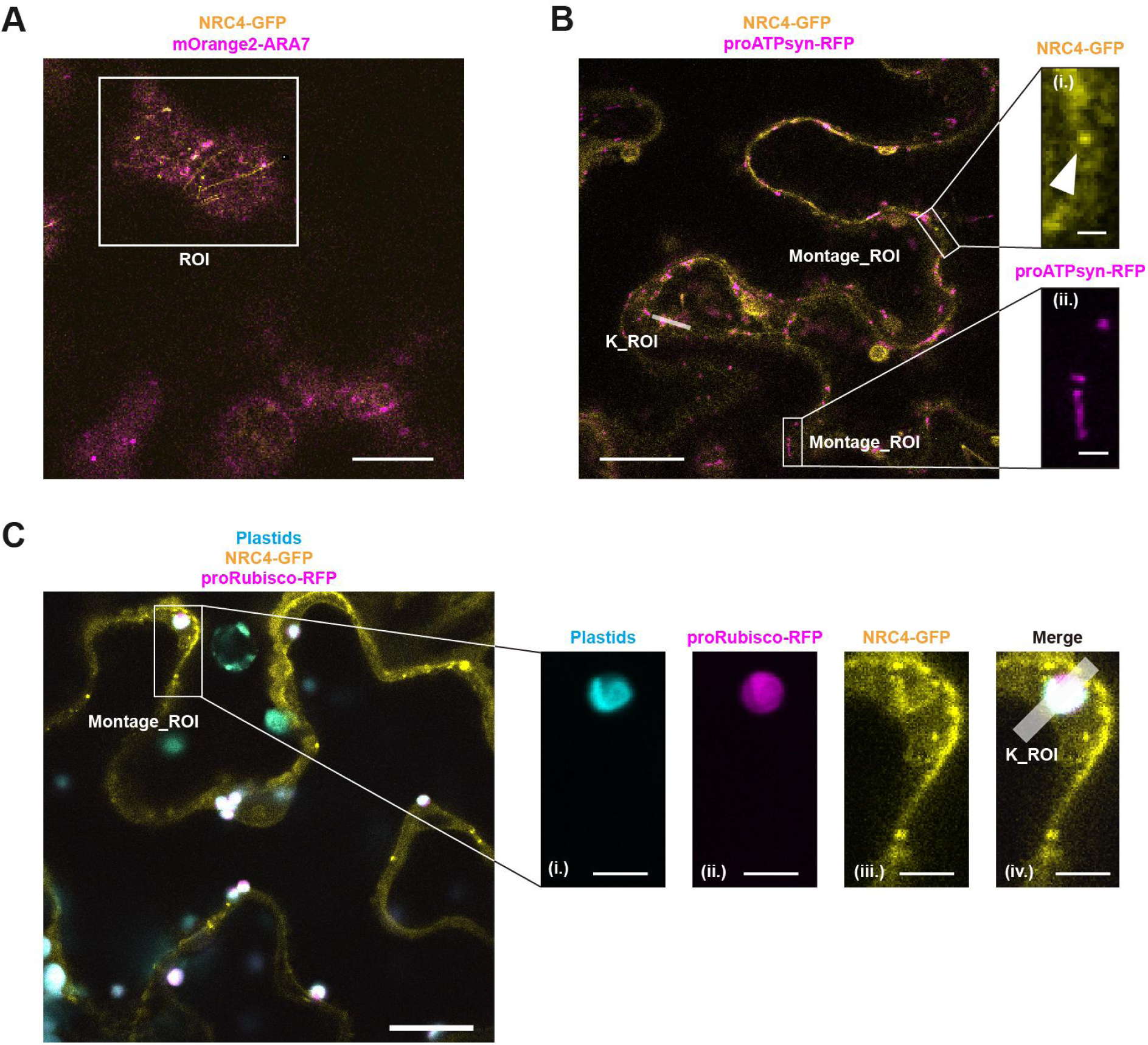
Regions of interest selected for generating montages and kymographs in Figure 4. A. Representative images showing overlaid NRC4-GFP/mOrange2-ARA7 corresponding with Movie S7. The white box indicates the region of interest (ROI) used to generate the time-lapse montages and for endosome-tracking shown in Fig. 4A and 4B. Scale bar = 10 μm. B. and C. Representative images showing NRC4-GFP overlaid with proATPsyn-RFP (B, corresponding to Movie S9) or with plastids and proRubisco-RFP (C, corresponding to Movie S10). Left panels in (B) and (C) show the original fields of view with white boxes marking the ROIs used for montages in Fig. 4C and 4F. Scale bars = 20 μm. Right panels in (B) and (C) display zoomed-in single-channel views. In (B), panels (i) and (ii) show NRC4-GFP (i) and proATPsyn-RFP (ii), with arrowheads indicating an NRC4-GFP punctum. Scale bar = 2 μm. In (C), panels (i)–(iv) show plastid autofluorescence (i), proRubisco-RFP (ii), NRC4-GFP (iii), and the merged image (iv). The white line denotes the ROI used for generating the kymograph (K_ROI) shown in Figure 4G. Scale bar = 5 μm.

**Figure S7.**
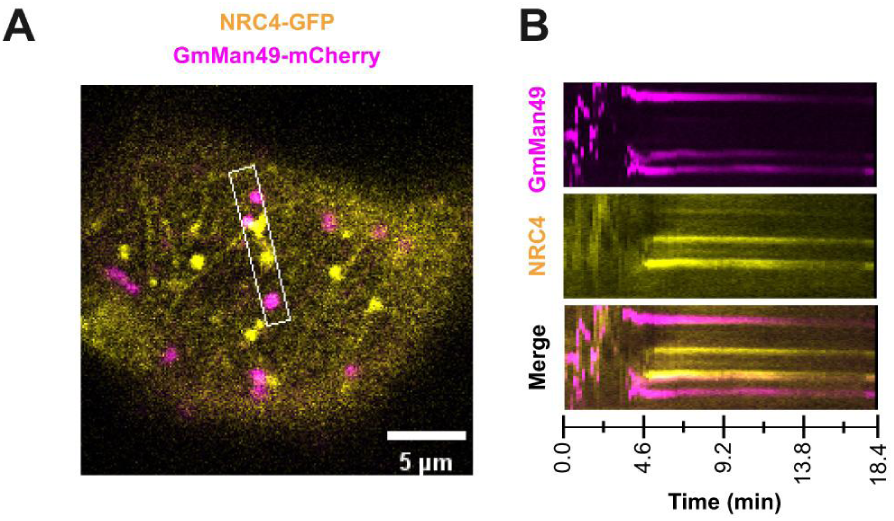
Cis-Golgi network movement ceases during hypersensitive cell death. A. Representative image showing region of interest (ROI) used for kymograph analysis in Fig. S4B. White area indicates the ROI selected for the kymograph. (see Movies S8.) B. Kymograph analysis of golgi dynamics during hypersensitive cell death. The top, middle and bottom panels show the signals of GmMan49-mCherry, NRC4-GFP and merged channel images, respectively. S10.)

**Figure S8.**
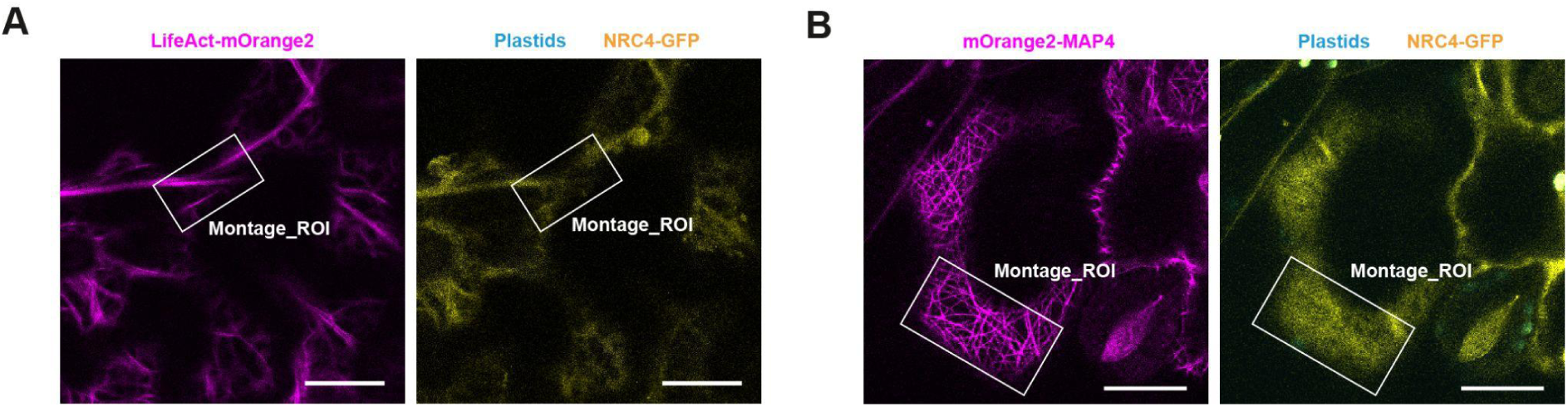
Regions of interest selected for generating montages and analyzing anisotropy in Figure 5. A. and B. Representative images showing LifeAct-mOrange2 (A, left) or mOrange2-MAP4 (B, left), and overlaid plastid/NRC4-GFP channels (A and B, right). The white boxes indicate the regions of interest (ROIs) used for montage generation and anisotropy quantification shown in Fig. 5A, 5B (for S8A), and 5C, 5D (for S8B). Scale bars = 20 μm.

**Figure S9.**
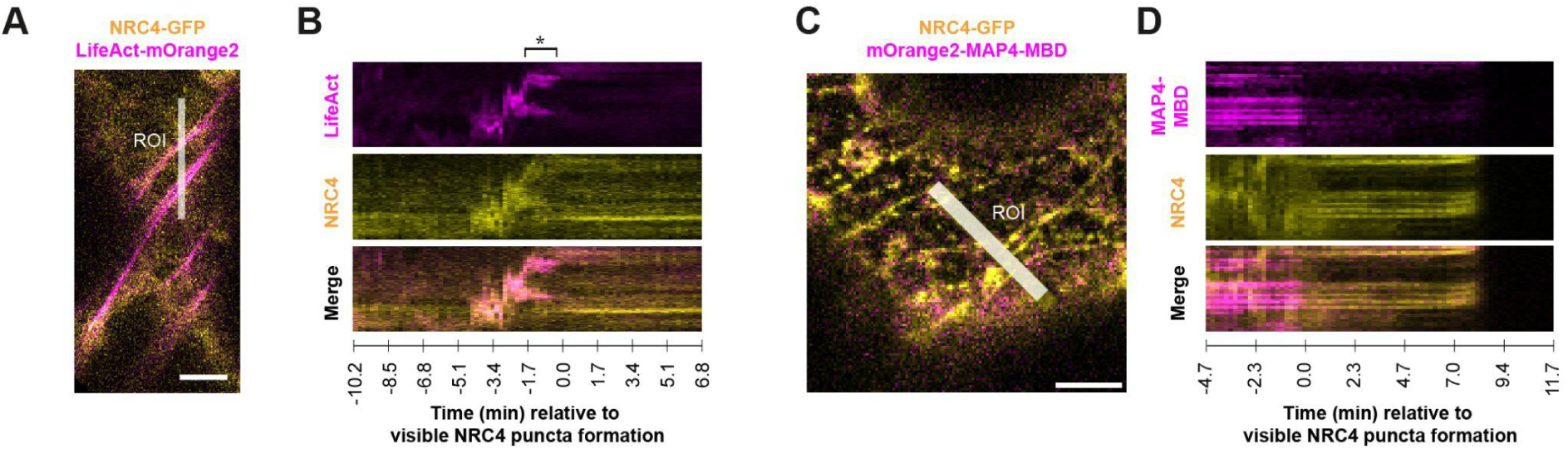
Actin movement stopped, and both cytoskeleton depolymerized during hypersensitive cell death. A. Merged image displaying the fluorescence signals of LifeAct-mOrange2-labeled actin and NRC4-GFP at the cell periphery. The white line marks the ROI used for kymograph analysis of both actin and NRC4 dynamics. Scale bar = 5 μm. (Movie S11) B. Kymographs showing the temporal dynamics of actin and NRC4 during hypersensitive cell death. The top, middle and bottom panels show the signals of LifeAct-mOrange2, NRC4-GFP, and composite image, respectively. The asterisk denotes the cessation of actin movement. C. Merged image displaying the fluorescence signals of mOrange2-MAP4-MBD-labeled microtubule and NRC4-GFP at the cell periphery. The white line marks the region of interest used for kymograph analysis of both microtubule and NRC4 dynamics. Scale bar = 5 μm. (Movie S13) D. Kymographs showing the temporal dynamics of microtubule and NRC4 dynamics during hypersensitive cell death. The top, middle and bottom panels show the signals of mOrange2-MAP4-MBD, NRC4-GFP, and composite image, respectively.

**Figure S10.**
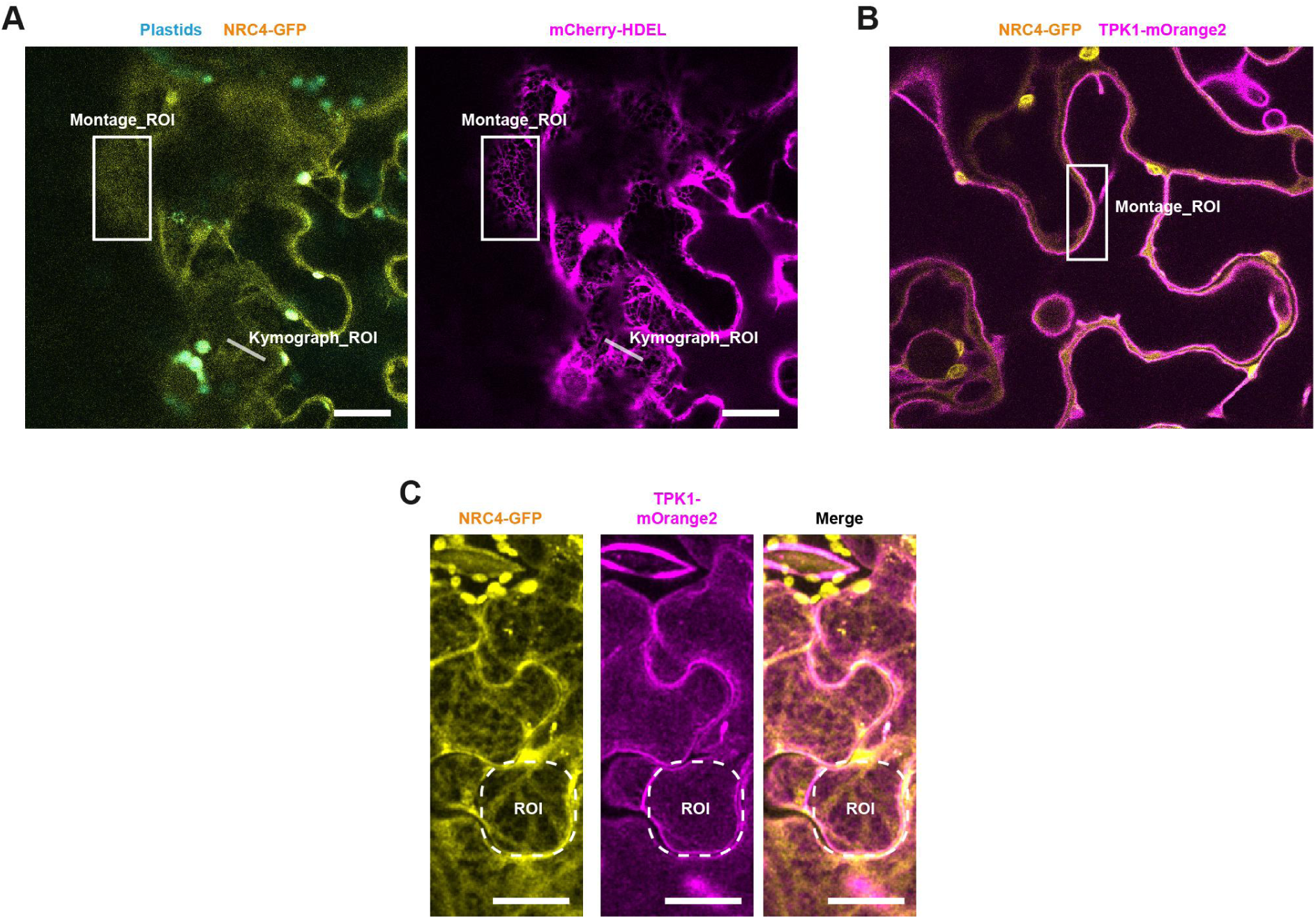
Regions of interest selected for analyzing endoplasmic reticulum (ER) and tonoplast. A. Representative images showing overlaid NRC4-GFP/plastids (left) and mCherry-HDEL (right) corresponding with Movie S14. The white box indicates the region of interest (ROI) used to generate the time-lapse montages in Fig. 6A. The white solid line indicates the ROI used for generating kymographs. Scale bar = 20 μm. B. Representative images showing overlaid NRC4-GFP/TPK1-mOrange2 corresponding with Movie S17. The white solid box indicates the ROI used to generate the time-lapse montages in Fig. 6F. Scale bar = 20 μm. C. Representative NRC4-GFP, TPK1-mOrange2 and merged channel images. The white circle indicates the ROI used for quantifying coverage of TPK1-mOrange2 in Fig. S11D. Scale bar = 20 μm.

**Figure S11.**
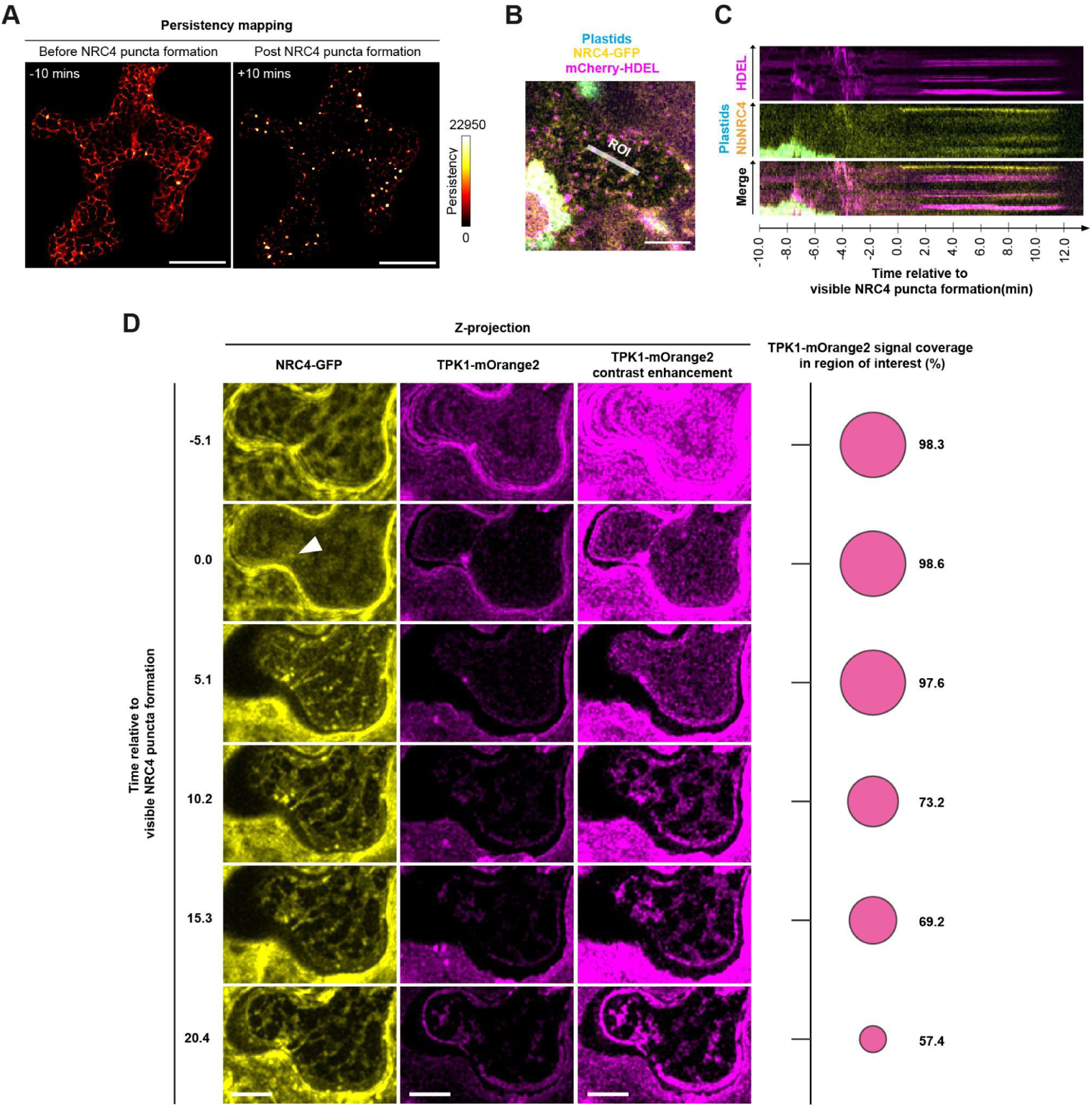
NRC4 resistosome activation affects endoplasmic reticulum (ER) tubule stability and tonoplast dynamic. A. Persistence mapping of ER structure before (left) and after (right) the formation of NRC4 puncta. A 10 minutes time-series of ER dynamics during hypersensitive cell death was segmented and processed to generate a substack of non-moving signals across time. Persistence maps were generated by a Z-projection of this non-moving substack, where the pixel value indicates signal persistence over time. Scale bar = 20 μm. B. Representative image showed the fragmented ER with NRC4 puncta. The white line marks the ROI used for kymograph analysis. Scale bar = 10 μm. (Movie S14) C. Kymograph analysis of ER dynamics during hypersensitive cell death. The top, middle and bottom panels show the signals of mCherry-HDEL, NRC4-GFP with plastid and merged channel images, respectively. D. Representative images of tonoplast membrane dynamics during hypersensitive cell death. The representative images were subset from the time series imaging with 5.1-minute intervals. Left panels: z-projection of NRC4-GFP, original and contrast-enhancing TPK1-mOrange2. The white arrowhead indicates an NRC4 punctum form on the cell periphery. Right panels: fluorescence coverage analysis of TPK1-mOrange2. Coverage (%) represents the area of TPK1-mOrange2, divided by the total area of ROI. Time zero is set relative to the onset of NRC4 puncta formation. Scale bar = 10 μm. (Movie S18)

**Figure S12.**
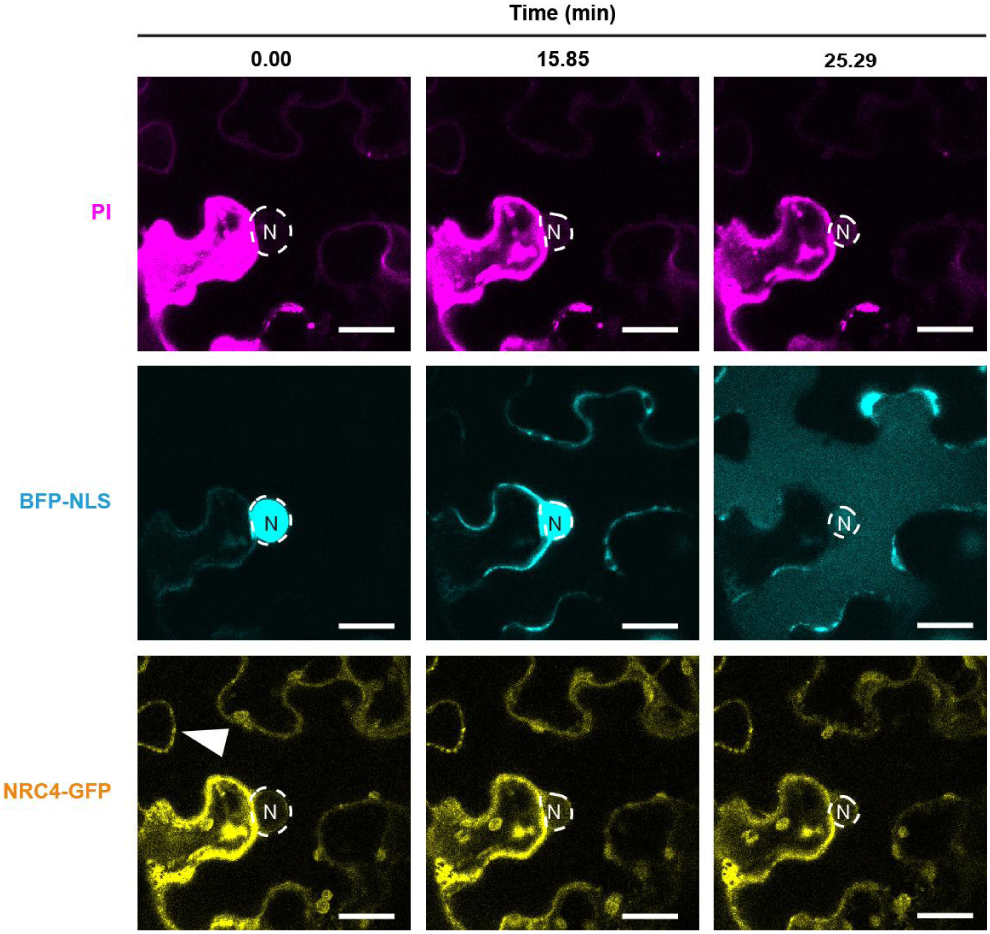
Nucleus and tonoplast disintegrate during hypersensitive cell death. Representative confocal images showing sequential events in a single cell: NRC4 puncta formation, BFP-NLS leakage, and tonoplast breakdown at 0.00, 15.85, and 25.29 minutes, respectively. Timepoints correspond to Movie S21. White circles indicate the nucleus area (N) in each panel. Top: propidium iodide; middle: BFP-NLS; bottom: NRC4-GFP. White arrowhead indicates an NRC4-GFP punctum. Scale bar = 20 μm.

**Figure S13.**
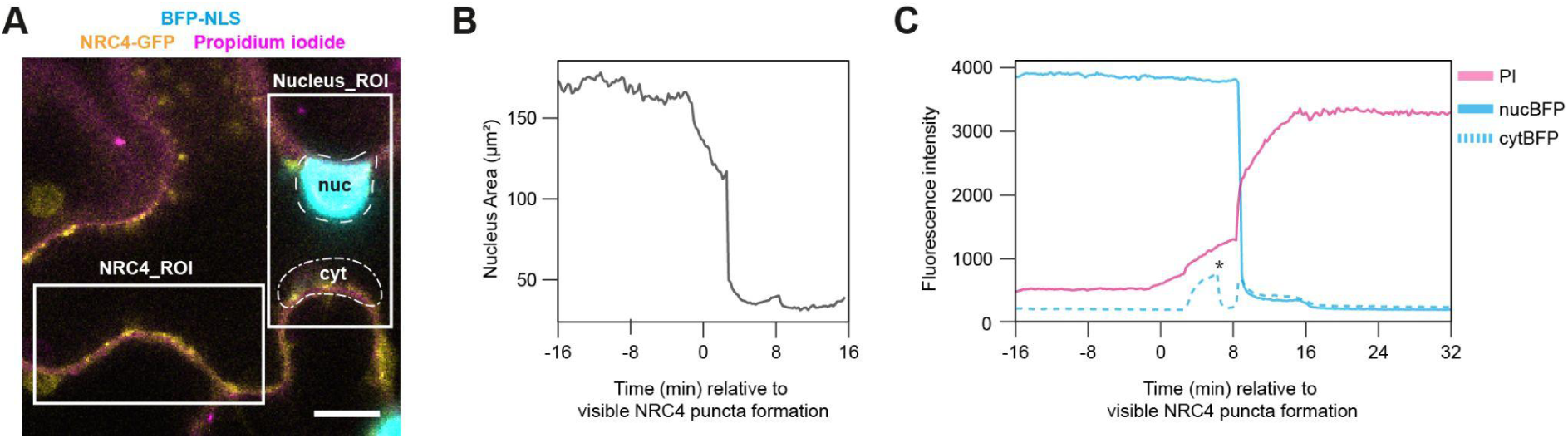
Nuclear shrinkage was followed by increased permeability to propidium iodide during cell death progression. A. Region of interests (ROIs) selected for quantification. White squares denote the areas selected for NRC4-GFP and the nuclear region used in Fig. 7A. The dashed line marks the nuclear region analyzed in Fig. S13B and S13C. The dash-dot line outlines the cytosolic region with BFP signal used in Fig. S13C. Scale bar = 10 μm. (see Movie S20) B. and C. Quantification of temporal changes in nuclear area (B) and fluorescence intensity of BFP-NLS and PI staining (C) during hypersensitive cell death. Nuclear region was selected for measuring PI (solid pink) and nucBFP (solid cyan) intensities, while the cytosol region was used for cytBFP (dash cyan). Asterisk in C marks the timing of a local tonoplast breakdown leading to decrease of cytosolic BFP intensity. Data corresponds to the cell shown in Figure 7A and Movie S20.

**Figure S14.**
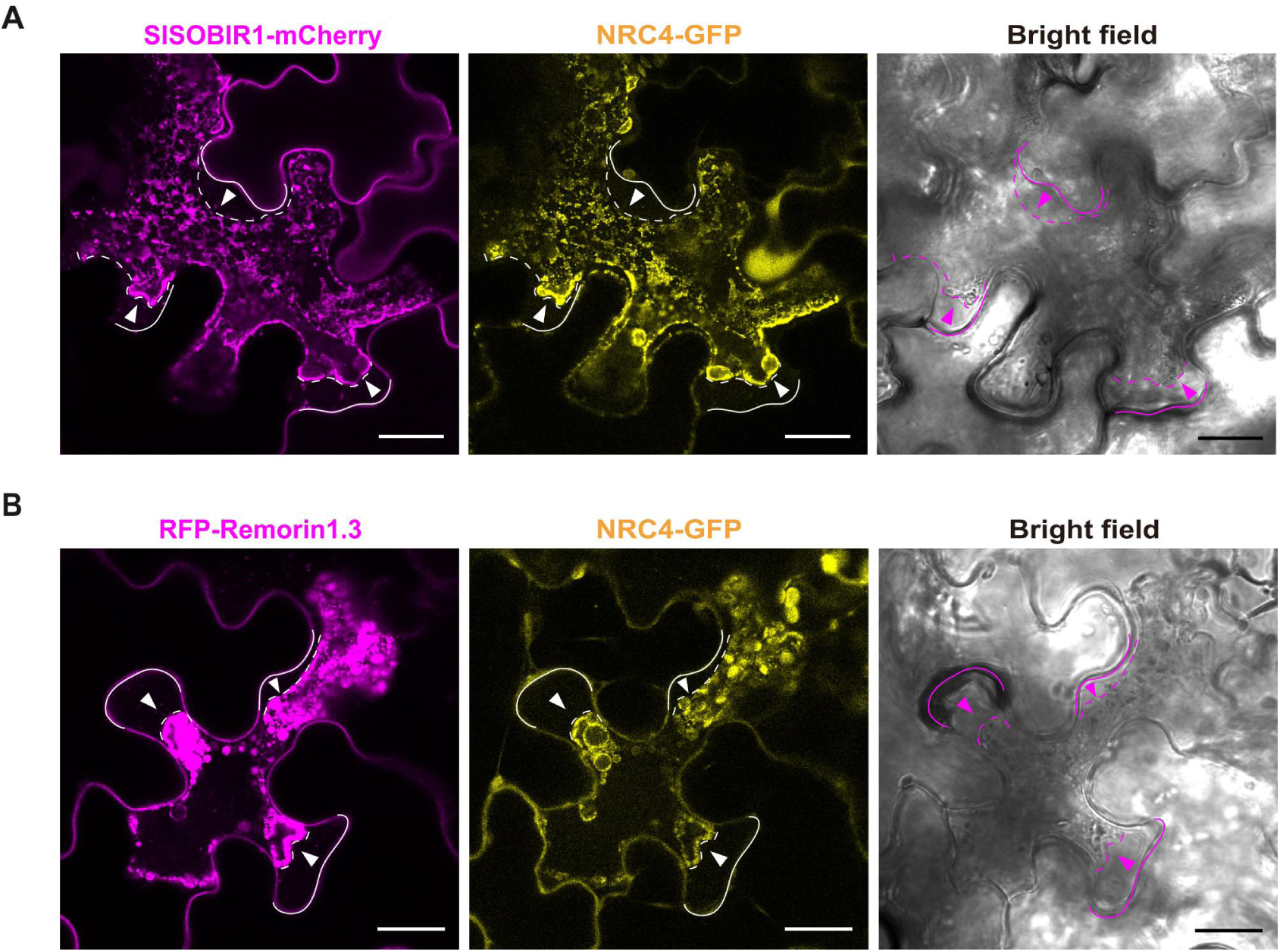
Plasma membrane shrank during hypersensitive cell death. A. and B. Representative confocal images showing the SlSOBI1 (A) or RFP-Remorin1.3 (B) labeled plasma membrane, NRC4-GFP (middle) and bright field (right). Solid and dotted lines mark the cell boundaries of the central and neighboring cells, respectively. Arrowheads mark regions where plasma membrane shrinkage is observed. Scale bar = 20 μm.

**Figure S15.**
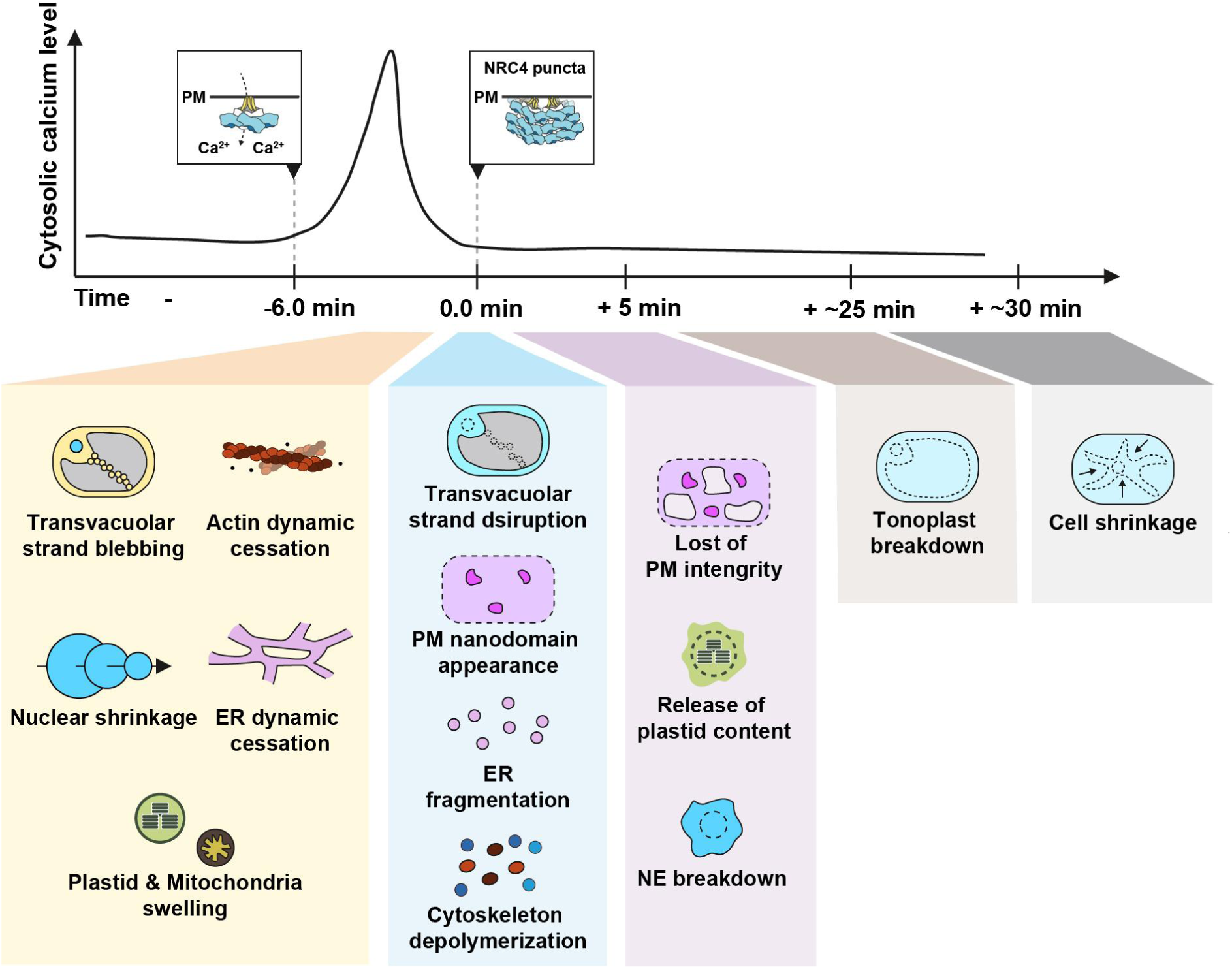
Dynamics of calcium and organelles in NRC4 resistosome mediated hypersensitive cell death. Activation of NRC4 triggers a transient influx of calcium into the cytosol. This calcium spike coincides with a halt in endoplasmic reticulum (ER) and actin dynamics, nuclear shrinkage, and swelling of plastids and mitochondria. As cell death progresses, the membrane systems become severely disrupted, with the appearance of plasma membrane (PM) nanodomains and the fragmentation of the ER, nuclear envelope (NE), and plastids. The cytoskeleton also undergoes depolymerization. At later stages, the tonoplast disintegrates along with a loss of PM integrity. Ultimately, the cell undergoes shrinkage, with both the PM and tonoplast collapsing inward toward the cell center.

**Movie S1.** NRC4^3A^-GFP forms puncta upon activation, related to Figure 2A, 2C, S2A and S3A.

**Movie S2.** NRC4^3A^-GFP relocalizes from cytosol to cell periphery upon activation, related to Figure S3B.

**Movie S3.** NRC4-GFP forms puncta and filamentous structures upon activation, followed by cell collapse. Related to Figures 2B, 2D-E, S2B and S3A.

**Movie S4.** NRC4-GFP relocalizes from cytosol to cell periphery upon activation. Related to Figure S3E.

**Movie S5.** A cytosolic calcium surge precedes cell collapse. Related to Figure 3A-B.

**Movie S6.** Enrichment of NRC4-GFP at the cell periphery coincides with the cytosolic calcium surge, followed by the formation of NRC4 puncta. Related to Figure 3C-D.

**Movie S7.** Endosomes labeled with ARA7-RFP stop trafficking prior to the formation of NRC4 puncta. Related to Figures 4A-B, and S6A.

**Movie S8.** Cis-Golgi cisternae labeled with GmMan49-RFP stop trafficking prior to NRC4 puncta formation. Related to Figures S7A-B.

**Movie S9.** Mitochondria labeled with proATPsyn-RFP stop trafficking prior to NRC4 puncta formation and transition into spherical, vesicle-like structures. Related to Figures 4C-4E and S6B.

**Movie S10.** Plastids labeled with proRubisco-RFP stop trafficking prior to NRC4 puncta formation, then swell and burst, releasing stromal contents into the cytosol after puncta formation. Related to Figures 4F-H, and S6C.

**Movie S11.** Actin dynamics labeled with LifeAct-mOrange2 stop prior to NRC4 puncta formation, with a marked decrease in signal coinciding with the onset of visible NRC4 puncta. Related to Figures 5A-B, S8A, and S9A-B.

**Movie S12.** Microtubules labeled with mOrange2-MAP4-MBD undergo depolymerization at the onset of NRC4 puncta formation. Related to Figures 5C-D, S8B, and S9C-D.

**Movie S13.** Activated NRC4 forms filamentous structures that align with microtubule patterns. Related to Figure S9C-D.

**Movie S14.** Endoplasmic reticulum labeled with mCherry-HDEL stops streaming prior to NRC4 puncta formation, then fragments into vesicle-like structures upon puncta formation. Related to Figure 6A, S10A, S11B-C.

**Movie S15.** Plasma membrane labeled with *Sl*SOBIR1-mCherry redistributes upon NRC4 puncta formation, and the signal decreases after cell collapse as the cell periphery moves out of the focal plane. Related to Figure 6C.

**Movie S16.** Plasma membrane lipids stained with FM4-64 remain stable before and after NRC4 puncta formation, with signal decrease after cell collapse as the cell periphery moves out of the focal plane. Related to Figure 6D.

**Movie S17.** Tonoplast labeled by *At*TPK1-mOrange2 lost integrity after NRC4 puncta formation. Related to Figure 6F, and S10B.

**Movie S18.** Reconstructed 3D time-lapse imaging of tonoplast membrane integrity during hypersensitive cell death. The tonoplast loses integrity after NRC4 puncta formation. Related to Figure S10C, and S11D.

**Movie S19.** Reconstructed 3D time-lapse imaging of tonoplast dynamics during hypersensitive cell death. Transvacuolar strands undergo blebbing and break down into vesicle-like structures during the process. Related to Figure 6G.

**Movie S20.** Nucleus labeled with BFP-NLS undergoes movement cessation, nuclear shrinkage, and propidium iodide (PI) entry during hypersensitive cell death. Related to Figure 7A-B, and S13A-C.

**Movie S21.** Representative time-lapse series showing a progression of events during hypersensitive cell death: nuclear shrinkage, nuclear envelope (NE) disintegration indicated by release of BFP-NLS into the cytosol, tonoplast (TN) disintegration indicated by entry of BFP-NLS into the vacuole, and propidium iodide (PI) entry. Related to Figure 7B, and S12.

**Movie S22.** Nuclear envelope labeled with AtPNET2 disperses into the cytosol after NRC4 puncta formation. Related to Figure 7C.

**Movie S23.** Tonoplast labeled with AtTPK1 detaches from the cell boundary at a late stage of cell death. Related to Figure 7D.

## References

Ahn H, Lin X, Olave-Achury AC, Derevnina L, Contreras MP, Kourelis J, Wu C, Kamoun S, and Jones JDG. Effector-dependent activation and oligomerization of plant NRC class helper NLRs by sensor NLR immune receptors Rpi-amr3 and Rpi-amr1. The EMBO Journal. 2023:42(5):e111484. 10.15252/embj.2022111484

Akerboom J, Carreras Calderón N, Tian L, Wabnig S, Prigge M, Tolö J, Gordus A, Orger MB, Severi KE, Macklin JJ, et al. Genetically encoded calcium indicators for multi-color neural activity imaging and combination with optogenetics. Front Mol Neurosci. 2013:6:2. 10.3389/fnmol.2013.00002

Burke R, McCabe A, Sonawane NR, Rathod MH, Whelan CV, McCabe PF, and Kacprzyk J. Arabidopsis cell suspension culture and RNA sequencing reveal regulatory networks underlying plant-programmed cell death. The Plant Journal. 2023:115(6):1465–1485. 10.1111/tpj.16407

Cai G, Parrotta L, and Cresti M. Organelle trafficking, the cytoskeleton, and pollen tube growth. Journal of Integrative Plant Biology. 2015:57(1):63–78. 10.1111/jipb.12289

Chai J, Song W, and Parker JE. New Biochemical Principles for NLR Immunity in Plants. MPMI. 2023:36(8):468–475. 10.1094/MPMI-05-23-0073-HH

Chen T-W, Wardill TJ, Sun Y, Pulver SR, Renninger SL, Baohan A, Schreiter ER, Kerr RA, Orger MB, Jayaraman V, et al. Ultrasensitive fluorescent proteins for imaging neuronal activity. Nature. 2013:499(7458):295–300. 10.1038/nature12354

Chiang B-J, Lin K-Y, Chen Y-F, Huang C-Y, Goh F-J, Huang L-T, Chen L-H, and Wu C-H. Development of a tightly regulated copper-inducible transient gene expression system in Nicotiana benthamiana incorporating a suicide exon and Cre recombinase. New Phytol. 2024:244(1):318–331. 10.1111/nph.20021

Coll NS, Epple P, and Dangl JL. Programmed cell death in the plant immune system. Cell Death Differ. 2011:18(8):1247–1256. 10.1038/cdd.2011.37

Coll NS, Smidler A, Puigvert M, Popa C, Valls M, and Dangl JL. The plant metacaspase AtMC1 in pathogen-triggered programmed cell death and aging: functional linkage with autophagy. Cell Death Differ. 2014:21(9):1399–1408. 10.1038/cdd.2014.50

Contreras MP, Pai H, Tumtas Y, Duggan C, Yuen ELH, Cruces AV, Kourelis J, Ahn H, Lee K, Wu C, et al. Sensor NLR immune proteins activate oligomerization of their NRC helpers in response to plant pathogens. The EMBO Journal. 2023:42(5):e111519. 10.15252/embj.2022111519

Du J, Verzaux E, Chaparro-Garcia A, Bijsterbosch G, Keizer LCP, Zhou J, Liebrand TWH, Xie C, Govers F, Robatzek S, et al. Elicitin recognition confers enhanced resistance to Phytophthora infestans in potato. Nat Plants. 2015:1(4):15034. 10.1038/nplants.2015.34

Duggan C, Moratto E, Savage Z, Hamilton E, Adachi H, Wu C-H, Leary AY, Tumtas Y, Rothery SM, Maqbool A, et al. Dynamic localization of a helper NLR at the plant–pathogen interface underpins pathogen recognition. Proceedings of the National Academy of Sciences. 2021:118(34):e2104997118. 10.1073/pnas.2104997118

Duxbury Z, Wu C, and Ding P. A Comparative Overview of the Intracellular Guardians of Plants and Animals: NLRs in Innate Immunity and Beyond. Annual Review of Plant Biology. 2021:72(Volume 72, 2021):155–184. 10.1146/annurev-arplant-080620-104948

Hofius D, Li L, Hafrén A, and Coll NS. Autophagy as an emerging arena for plant–pathogen interactions. Current Opinion in Plant Biology. 2017:38:117–123. 10.1016/j.pbi.2017.04.017

Hofius D, Schultz-Larsen T, Joensen J, Tsitsigiannis DI, Petersen NHT, Mattsson O, Jørgensen LB, Jones JDG, Mundy J, and Petersen M. Autophagic components contribute to hypersensitive cell death in Arabidopsis. Cell. 2009:137(4):773–783. 10.1016/j.cell.2009.02.036

Huang S, Li E, Jia F, Han Z, and Chai J. Assembly and functional mechanisms of plant NLR resistosomes. Current Opinion in Structural Biology. 2025a:90:102977. 10.1016/j.sbi.2024.102977

Huang S, Wang J, Song R, Jia A, Xiao Y, Sun Y, Wang L, Mahr D, Wu Z, Han Z, et al. Balanced plant helper NLR activation by a modified host protein complex. Nature. 2025b:639(8054):447–455. 10.1038/s41586-024-08521-7

Kasaras A and Kunze R. Dual-targeting of Arabidopsis DMP1 isoforms to the tonoplast and the plasma membrane. PLOS ONE. 2017:12(4):e0174062. 10.1371/journal.pone.0174062

Kourelis J, Contreras MP, Harant A, Pai H, Ludke D, Adachi H, Derevnina L, Wu CH, and Kamoun S. The helper NLR immune protein NRC3 mediates the hypersensitive cell death caused by the cell-surface receptor Cf-4. PLoS Genet. 2022:18(9):e1010414. 10.1371/journal.pgen.1010414

Lee S, Lee DW, Yoo Y-J, Duncan O, Oh YJ, Lee YJ, Lee G, Whelan J, and Hwang I. Mitochondrial Targeting of the Arabidopsis F1-ATPase γ-Subunit via Multiple Compensatory and Synergistic Presequence Motifs. The Plant Cell. 2012:24(12):5037–5057. 10.1105/tpc.112.105361

Liu F, Yang Z, Wang C, You Z, Martin R, Qiao W, Huang J, Jacob P, Dangl JL, Carette JE, et al. Activation of the helper NRC4 immune receptor forms a hexameric resistosome. Cell. 2024:187(18):4877–4889.e15. 10.1016/j.cell.2024.07.013

Ma S, An C, Lawson AW, Cao Y, Sun Y, Tan EYJ, Pan J, Jirschitzka J, Kümmel F, Mukhi N, et al. Oligomerization-mediated autoinhibition and cofactor binding of a plant NLR. Nature. 2024:632(8026):869–876. 10.1038/s41586-024-07668-7

Madhuprakash J, Toghani A, Contreras MP, Posbeyikian A, Richardson J, Kourelis J, Bozkurt TO, Webster MW, and Kamoun S. A disease resistance protein triggers oligomerization of its NLR helper into a hexameric resistosome to mediate innate immunity. Sci Adv. 2024:10(45):eadr2594. 10.1126/sciadv.adr2594

Madina MH, Rahman MS, Zheng H, and Germain H. Vacuolar membrane structures and their roles in plant–pathogen interactions. Plant Mol Biol. 2019:101(4):343–354. 10.1007/s11103-019-00921-y

Maekawa T, Kashkar H, and Coll NS. Dying in self-defence: a comparative overview of immunogenic cell death signalling in animals and plants. Cell Death Differ. 2023:30(2):258–268. 10.1038/s41418-022-01060-6

Mur LAJ, Kenton P, Lloyd AJ, Ougham H, and Prats E. The hypersensitive response; the centenary is upon us but how much do we know? J Exp Bot. 2008:59(3):501–520. 10.1093/jxb/erm239

Nelson BK, Cai X, and Nebenführ A. A multicolored set of in vivo organelle markers for co-localization studies in Arabidopsis and other plants. The Plant Journal. 2007:51(6):1126–1136. 10.1111/j.1365-313X.2007.03212.x

Pain C, Tolmie F, Wojcik S, Wang P, and Kriechbaumer V. intER-ACTINg: The structure and dynamics of ER and actin are interlinked. Journal of Microscopy. 2023:291(1):105–118. 10.1111/jmi.13139

Peng K-C, Wang C-W, Wu C-H, Huang C-T, and Liou R-F. Tomato SOBIR1/EVR Homologs Are Involved in Elicitin Perception and Plant Defense Against the Oomycete Pathogen Phytophthora parasitica. MPMI. 2015:28(8):913–926. 10.1094/MPMI-12-14-0405-R

Perico C and Sparkes I. Plant organelle dynamics: cytoskeletal control and membrane contact sites. New Phytologist. 2018:220(2):381–394. 10.1111/nph.15365

Salguero-Linares J, Serrano I, Ruiz-Solani N, Salas-Gómez M, Phukan UJ, González VM, Bernardo-Faura M, Valls M, Rengel D, and Coll NS. Robust transcriptional indicators of immune cell death revealed by spatiotemporal transcriptome analyses. Molecular Plant. 2022:15(6):1059–1075. 10.1016/j.molp.2022.04.010

Scheuring D, Viotti C, Krüger F, Künzl F, Sturm S, Bubeck J, Hillmer S, Frigerio L, Robinson DG, Pimpl P, et al. Multivesicular Bodies Mature from the Trans-Golgi Network/Early Endosome in Arabidopsis[W]. Plant Cell. 2011:23(9):3463–3481. 10.1105/tpc.111.086918

Selvaraj M, Toghani A, Pai H, Sugihara Y, Kourelis J, Yuen ELH, Ibrahim T, Zhao H, Xie R, Maqbool A, et al. Activation of plant immunity through conversion of a helper NLR homodimer into a resistosome. PLOS Biology. 2024:22(10):e3002868. 10.1371/journal.pbio.3002868

Shen Q, Hasegawa K, Oelerich N, Prakken A, Tersch LW, Wang J, Reichhardt F, Tersch A, Choo JC, Timmers T, et al. Cytoplasmic calcium influx mediated by plant MLKLs confers TNL-triggered immunity. Cell Host & Microbe. 2024:32(4):453–465.e6. 10.1016/j.chom.2024.02.016

Sunil S, Beeh S, Stöbbe E, Fischer K, Wilhelm F, Meral A, Paris C, Teasdale L, Jiang Z, Zhang L, et al. Activation of an atypical plant NLR with an N-terminal deletion initiates cell death at the vacuole. EMBO Rep. 2024:25(10):4358–4386. 10.1038/s44319-024-00240-4

Tang Y, Dong Q, Wang T, Gong L, and Gu Y. PNET2 is a component of the plant nuclear lamina and is required for proper genome organization and activity. Developmental Cell. 2022:57(1):19–31.e6. 10.1016/j.devcel.2021.11.002

Thomas CM, Jones DA, Parniske M, Harrison K, Balint-Kurti PJ, Hatzixanthis K, and Jones JD. Characterization of the tomato Cf-4 gene for resistance to Cladosporium fulvum identifies sequences that determine recognitional specificity in Cf-4 and Cf-9. Plant Cell. 1997:9(12):2209–2224. 10.1105/tpc.9.12.2209

Wang H-Y, Yuen ELH, Lee K-T, Goh F-J, Bozkurt TO, and Wu C-H. A hydrophobic core in the coiled-coil domain is essential for NRC resistosome function. 2025:2025.01.21.634219. 10.1101/2025.01.21.634219

Wang J, Bollier N, Buono RA, Vahldick H, Lin Z, Feng Q, Hudecek R, Jiang Q, Mylle E, Van Damme D, et al. A developmentally controlled cellular decompartmentalization process executes programmed cell death in the Arabidopsis root cap. Plant Cell. 2023:36(4):941–962. 10.1093/plcell/koad308

Wang P and Hussey PJ. Interactions between plant endomembrane systems and the actin cytoskeleton. Front Plant Sci. 2015:6. 10.3389/fpls.2015.00422

Wu C-H, Abd-El-Haliem A, Bozkurt TO, Belhaj K, Terauchi R, Vossen JH, and Kamoun S. NLR network mediates immunity to diverse plant pathogens. Proceedings of the National Academy of Sciences. 2017:114(30):8113–8118. 10.1073/pnas.1702041114

Yu H, Xu W, Chen S, Wu X, Rao W, Liu X, Xu X, Chen J, Nishimura MT, Zhang Y, et al. Activation of a helper NLR by plant and bacterial TIR immune signaling. Science. 2024:386(6728):1413–1420. 10.1126/science.adr3150

