## Supplemental materials for "Single-cell-resolved calcium and organelle dynamics in resistosome-mediated cell death"

### Materials and Methods

Plant growth conditions

Wild-type, *nrc2/3/4* knockout, and GCaMP6-expressing *Nicotiana benthamiana* plants were used for the analyses (Wu et al. 2020). Plants were cultivated in a walk-in growth chamber under a 16-hour light/8-hour dark photoperiod, with temperature maintained at 25 °C and relative humidity set between 45% and 65%.

Plasmid construction

Constructs used in this study were generated using Golden Gate assembly following the MoClo system (Weber et al. 2011; Engler et al. 2014). Level 0 plasmids of the copper-inducible system (CUP2, p65, CBS4-CMV1, CBS4-miniDFR, LoxP-mCherry-stop, and Cre) were described previously (Chiang et al. 2024). Level 0 modules of the RCaMP1h, proATPsyn, and AtPNET2 were synthesized as MoClo-compatible modules in kanamycin-resistant pUC57 vectors provided by SynBio Technologies (New Jersey, USA). For the cloning of GCaMP6, MAP4-MBD, ARA7, and LifeAct, sequences were amplified from plasmids kindly provided by Dr. Yen-Ping Hsueh (Institute of Molecular Biology, Academia Sinica) and Dr. Tien-Shin Yu (Institute of Plant Microbial Biology, Academia Sinica). *AtTPK1* was amplified from *Arabidopsis thaliana* cDNA. For fluorescent protein tags, mCherry2 was amplified from the plasmid pEB2-mCherry-L (Addgene #104002) (Balleza et al. 2018), mOrange2 was amplified from plasmid pBI-OFP (Lee et al. 2022), and mTagBFP2 was amplified from plasmid pICSL50021 provided by Mark Youles (SynBio, The Sainsbury Laboratory, UK). Sequences containing internal BsaI or BpiI sites were domesticated to be MoClo-compatible. All amplified sequences were cloned into pAGM9121 to generate MoClo-compatible Level 0 modules. The NRC4 and NRC2 variants were cloned into pICH86988, a binary vector that carries the 35S promoter and OCS terminator. The other modules were assembled into level 1 acceptors with different positions (pICH47751, pICH47761, pICH47781, and pICH47791) as indicated in Data S2, with 35S or CBS4 promoters, and NOS or OCS terminators. Sequences of plasmids, biosensors, subcellular markers, NLRs, AVRs, and primers used in this study are listed in Data S1 and S2.

Agroinfiltration and copper-induced gene expression

*Agrobacterium tumefaciens* strain GV3101/pMP90 was used for transient gene expression in this study. Bacterial strains were revived from glycerol stocks and cultured on 523 agar plates supplemented with appropriate antibiotics. After overnight incubation at 28 °C, cells were collected by centrifugation at 5,000 × g for 5 minutes at room temperature. The bacterial pellet was resuspended in MMA buffer (10 mM $MgCl_2$, 10 mM MES-KOH, 150 µM acetosyringone, pH 5.6) to an $OD_{600}$ as indicated in Table S1 and infiltrated into the 5$^{th}$ and 6$^{th}$ leaves of 4-week-old *N. benthamiana* plants using 1 mL syringes. For copper-induced gene expression, 10 µM $CuSO_4$ was infiltrated at the agroinfiltrated sites using 1 mL syringes at 2 days post-agroinfiltration.

<u>Cell death quantification</u>

For the whole leaf cell death assay, cell death was quantified using UVP ChemStudio Imaging Systems (Analytik Jena) as described previously at 36 hours post-copper infiltration (Goh et al. 2024). Raw cell death autofluorescence images were acquired using blue LED light for excitation and FITC filter (513-557 nm) as the emission filter. Areas with stronger visual cell death phenotype produce stronger autofluorescence signals. The exposure time was adjusted to 10 seconds to avoid saturation of the signal. The mean signal intensity was calculated using VisionWorks v.11.2 software by manually selecting the infiltrated areas and subtracting the background signal intensity. The mean signal intensity value was further normalized with the maximum intensity (65535) to obtain the relative intensity of cell death.

For the leaf disc assay, two days after agroinfiltration, the infiltrated areas were treated with either 10 µM $CuSO_4$ or $ddH_2O$. Leaf discs were collected using a 4 mm diameter cork borer and transferred to 96-well plates containing 200 µL of either 10 µM $CuSO_4$ or $ddH_2O$. Autofluorescence from was measured using a Synergy H1 plate reader (BioTek), with excitation and emission wavelengths set at 400 nm and 510 nm, respectively. Signals were recorded every 15 minutes over a 12-hour period, and data were acquired using Gen5 software (version 3.09, BioTek). Autofluorescence from copper-treated samples was normalized to the corresponding water-treated controls. Each experiment was performed with three biological replicates, each consisting of 11–12 leaf discs.

<u>Protein extraction and western blot analysis</u>

Protein extraction, SDS-PAGE electrophoresis, and Western blot analysis were conducted as previously described with minor modifications (Win et al. 2011). At the indicated time points following copper treatment, leaf discs were collected into 2 mL microtubes containing ceramic beads and immediately flash-frozen in liquid nitrogen. The frozen tissues were ground using an SH-100 homogenizer (J&H Technology Co., Ltd.) at 800 rpm for 30 seconds, repeated five times. Total protein was extracted using an extraction buffer containing 10% glycerol, 25 mM Tris (pH 7.5), 1 mM EDTA, 150 mM NaCl, 2% (w/v) PVPP, 10 mM DTT, 1× protease inhibitor cocktail (Sigma), and 0.1% IGEPAL (Sigma). The lysates were centrifuged at 13,000 × g for 10 minutes at 4 °C, and the supernatant was collected and mixed with 4× sample loading dye (200 mM Tris-HCl pH 6.8, 8% SDS, 40% glycerol, 50 mM EDTA, 0.08% bromophenol blue, and 100 mM DTT). Protein samples were denatured at 70 °C for 10 minutes and then separated on 10% or 15% polyacrylamide gels prepared with T-Pro EZ Gel Solution, 0.1% TEMED (Bio-Rad), and 0.1% ammonium persulfate (Bio-Rad). Following electrophoresis, proteins were transferred to PVDF membranes using the Trans-Blot Turbo Transfer System (Bio-Rad). Immunoblotting was performed using the SNAP i.d. 2.0 Protein Detection System (Merck) with anti-myc (A00704, GenScript) as the primary antibody and Peroxidase AffiniPure Goat Anti-Mouse IgG (H+L) (115-035-003, Jackson) as the secondary antibody. Chemiluminescent images were captured using the UVP ChemStudio Imaging System (Analytik Jena). SimplyBlue SafeStain (465034, Invitrogen) was used to stain the PVDF membrane to visualize rubisco.

Blue native PAGE (BN-PAGE)

The BN-PAGE analysis was conducted as previously described (Contreras et al. 2023). To detect the formation of NRC4 resistosome complexes, leaf discs were collected at the indicated time points following copper treatment, placed into 2 mL microtubes containing ceramic beads, and flash-frozen in liquid nitrogen. Tissues were ground using an SH-100 homogenizer (J&H Technology Co., Ltd.) at 800 rpm for 30 seconds, repeated five times. Total protein was extracted using a buffer containing 50 mM HEPES (pH 7.5), 50 mM NaCl, 5 mM $MgCl_2$, 10% glycerol, 10 mM DTT, 1× protease inhibitor cocktail (Sigma, P9599), and 1% digitonin (Sigma, D141). Extracts were incubated at 4 °C for 10 minutes and centrifuged twice at 13,000 × g for 10 minutes at 4 °C. The resulting supernatant was collected, diluted to 0.4× with extraction buffer, and mixed with 4× NativePAGE Sample Buffer (Invitrogen, BN2003) and NativePAGE 5% G-250 Sample Additive (Invitrogen, BN2004) to a final G-250 concentration of 0.125%. Proteins were separated on NativePAGE 4–16% Bis-Tris Gels (Invitrogen, BN1002). Following electrophoresis, proteins were transferred to PVDF membranes using the Trans-Blot Turbo Transfer System (Bio-Rad). Membranes were fixed in 8% acetic acid for 15 minutes, air-dried in a chemical fume hood, and rinsed with 99% ethanol until the blue dye was removed. Finally, membranes were rinsed with TBST and subjected to Western blot analysis as described above.

Time-lapse imaging of resistosomes, calcium, and organelles

Confocal laser scanning microscopy was conducted using an Olympus FV3000 confocal microscope equipped with a 40× silicone immersion objective, or a Leica STELLARIS 8 confocal microscope equipped with a 63× water immersion objective lens. At indicated time points after agroinfiltration, leaf discs were placed on Nunc Lab-Tek chambered coverglass (155361, Thermo Scientific) and were covered with a 2% water agar (agarose) to prevent dehydration during image acquisition. Time-lapse image acquisition was performed using Olympus FV3000 confocal microscope with FV31S-SW software (Evident Scientific) with a 3-5 hour imaging program. Fluorescence detection was set as the following excitation/emission settings: 405/445–480 nm for mTagBFP2, 488/505–540 nm for eGFP and GCaMP6, 561/565–620 nm for mOrange2, and 561/600–640 nm for mRFP, mCherry2, RCaMP1h, propidium iodide, and FM4-64. Subcellular markers and biosensors used in this study are listed in Table S1. For the 3D time-lapse experiments shown in Figures 6G, 7C, S10C, and 11D, *xyzt* scanning was performed.

PI and FM4-64 staining

For dead cell staining, leaves expressing the indicated constructs were infiltrated with 25 μg/mL propidium iodide (PI; P4864, Sigma-Aldrich). For plasma membrane staining, leaves were infiltrated with 10 μg/mL FM4-64 (T3166, Invitrogen). In experiments involving copper treatment, the chemical probes were co-applied with 10 μM copper solution at 30 minutes prior to imaging for proper staining.

Image analysis

Image analysis was performed by FIJI imageJ(1.54p) (Schindelin et al. 2012). Drift-correction for time-lapse imaging were performed using Fast4DReg plugin (Laine et al. 2019). Kymographs were generated using a Live-Kymographer plugin embedded in ImageJ. Colocalization analysis between NRC4 and cytosolic markers was quantified using the JaCoP plugin by calculating the Pearson's correlation coefficient between two fluorescence signals (Bolte and Cordelières 2006). Vesicle dynamics were analyzed using the TrackMate plugin with DoG detector for spot identification and the Simple LAP tracker for tracking (max linking distance: 5 μm; gap-closing distance: 4 μm; max frame gap: 2) (Ershov et al. 2022). Vesicle velocities were extracted from resulting tracks. For organelle roundness quantification, organelle areas were measured and analyzed using the Shape Descriptors function in ImageJ. Consistency mapping and mesh counting of ER were performed as previously described (Griffing 2018). For consistency mapping, time-lapse images of the ER were processed by image subtraction to generate a dataset representing non-mobile ER regions. The resulting image stack was used to calculate persistence values and visualized using the Red-Hot pseudocolor look-up table. For ER mesh counting, a binary mask of ER meshes was generated, and the number of meshes was quantified using the Analyze Particles function in ImageJ. Plasma membrane, and tonoplast coverage was measured by fluorescence area within a defined ROI and presented as percentage (%) of coverage.

Statistical analysis

Box and violin plots, and generalized linear model (GLM) were generated using R with the ggplot2 package (Wickham 2016). In box plots, boxes represent the interquartile range (IQR; 25th–75th percentiles), with the median indicated by a horizontal line. Whiskers extend to the most extreme values within 1.5× IQR from the quartiles. Violin plots depict the data distribution estimated by kernel density, where the width reflects local data density. Median values are marked by horizontal lines, and individual data points are overlaid with jitter to enhance visibility. Statistical comparisons of vesicle velocity across time points were conducted using the Kruskal-Wallis test, followed by Dunn's post hoc test for pairwise comparisons when significant differences were detected ($p < 0.05$). Groups labeled with different letters denote statistically significant differences.

**Table S1. List of constructs used for agroinfiltration assays.**

| Vector backbone | Promoter | Protein name | Tag | $OD_{600}$ | Reference |
|---|---|---|---|---|---|
| **NLRs, AVRs and regulatory elements** | | | | | |
| pICH86988 | 35S | NRC2 | C-terminal myc | 0.5 | This study |
| pICH86988 | 35S | $NRC4^{3A}$ | C-terminal myc | 0.5 | This study |
| pICH86988 | 35S | $NRC4^{L9E}$ | C-terminal myc | 0.5 | This study |
| pICH86988 | 35S | $NRC4^{K190R}$ | C-terminal myc | 0.5 | This study |
| pICH86988 | 35S | $NRC4^{3A}$ | C-terminal GFP | 0.5 | This study |
| pICH86988 | 35S | NRC4 | C-terminal GFP | 0.5 | This study |
| pICH47751 | 35S | Rpi-blb2 | none | 0.2 | (Chiang et al. 2024) |
| pICH47761 | CBS4 | AVRblb2 | none | 0.1 | (Chiang et al. 2024) |
| pICH47742 | 35S | CUP2 | C-terminal p65 | 0.2 | (Chiang et al. 2024) |
| pICH47761 | CBS4 | CP | N-terminal LoxP-mCherry | 0.1 | (Chiang et al. 2024) |
| pICH47791 | CBS4 | Cre | none | 0.2 | (Chiang et al. 2024) |
| **Subcellular markers and biosensors** | | | | | |
| pICH47781 | 35S | ARA7 | N-terminal mCherry | 0.1 | This study |
| pICH47781 | 35S | Lifeact | C-terminal mOrange2 | 0.1 | This study |
| pICH47781 | 35S | MAP4-MBD | N-terminal mOrange2 | 0.1 | This study |
| pICH47781 | 35S | AtTPK1 | C-terminal mOrange2 | 0.2 | This study |
| pICH47781 | 35S | AtPNET2 | C-terminal mOrange2 | 0.2 | This study |
| pICH47742 | 35S | GCaMP6 | N-terminal 6xHis | 0.1 | This study |
| pICH47781 | 35S | RCaMP1h | N-terminal 6xHis | 0.1 | This study |
| pICH47781 | 35S | mTagBFP2 | C-terminal NLS | 0.05 | This study |
| pGWB555 | 35S | mRFP | none | 0.1 | (Bozkurt et al. |

| | | | | | |
|---|---|---|---|---|---|
| | | | | | 2011) |
| | 35S | SlSOBIR1 | C-terminal mCherry | 0.1 | (Li et al. 2021) |
| | 35S | proATPsyn | C-terminal RFP | 0.1 | This study |
| | 35S | proRubisco | C-terminal RFP | 0.1 | (Nelson et al. 2007) |
| | 35S | GmMan49 | C-terminal RFP | 0.1 | (Nelson et al. 2007) |
| | 35S | mCherry | C-terminal HDEL | 0.1 | (Nelson et al. 2007) |

**Data S1. List of primers used in this study. (as a separate file)**

**Data S2. List of plasmids and sequences used in this study. (as a separate file)**

### References

**Balleza E, Kim JM, and Cluzel P**. Systematic characterization of maturation time of fluorescent proteins in living cells. Nat Methods. 2018:**15**(1):47–51. https://doi.org/10.1038/nmeth.4509

**Bolte S and Cordelières FP**. A guided tour into subcellular colocalization analysis in light microscopy. J Microsc. 2006:**224**(3):213–232. https://doi.org/10.1111/j.1365-2818.2006.01706.x

**Bozkurt TO, Schornack S, Win J, Shindo T, Ilyas M, Oliva R, Cano LM, Jones AME, Huitema E, van der Hoorn RAL, et al.** Phytophthora infestans effector AVRblb2 prevents secretion of a plant immune protease at the haustorial interface. Proc Natl Acad Sci. 2011:**108**(51):20832–20837. https://doi.org/10.1073/pnas.1112708109

**Chiang B-J, Lin K-Y, Chen Y-F, Huang C-Y, Goh F-J, Huang L-T, Chen L-H, and Wu C-H**. Development of a tightly regulated copper-inducible transient gene expression system in Nicotiana benthamiana incorporating a suicide exon and Cre recombinase. New Phytol. 2024:**244**(1):318–331. https://doi.org/10.1111/nph.20021

**Contreras MP, Pai H, Tumtas Y, Duggan C, Yuen ELH, Cruces AV, Kourelis J, Ahn H, Lee K, Wu C, et al.** Sensor NLR immune proteins activate oligomerization of their NRC helpers in response to plant pathogens. EMBO J. 2023:**42**(5):e111519. https://doi.org/10.15252/embj.2022111519

**Engler C, Youles M, Gruetzner R, Ehnert T-M, Werner S, Jones JDG, Patron NJ, and Marillonnet S**. A Golden Gate Modular Cloning Toolbox for Plants. ACS Synth Biol. 2014:**3**(11):839–843. https://doi.org/10.1021/sb4001504

**Ershov D, Phan M-S, Pylvänäinen JW, Rigaud SU, Le Blanc L, Charles-Orszag A, Conway JRW, Laine RF, Roy NH, Bonazzi D, et al.** TrackMate 7: integrating state-of-the-art segmentation algorithms into tracking pipelines. Nat Methods. 2022:**19**(7):829–832. https://doi.org/10.1038/s41592-022-01507-1

**Goh F-J, Huang C-Y, Derevnina L, and Wu C-H**. NRC Immune receptor networks show diversified hierarchical genetic architecture across plant lineages. Plant Cell. 2024:**36**(9):3399–3418. https://doi.org/10.1093/plcell/koae179

**Griffing LR**. Dancing with the Stars: Using Image Analysis to Study the Choreography of the Endoplasmic Reticulum and Its Partners and of Movement Within Its Tubules. . In. The Plant Endoplasmic Reticulum : Methods and Protocols, C Hawes and V Kriechbaumer, eds. (Springer: New York, NY), pp. 75–102. https://doi.org/10.1007/978-1-4939-7389-7_7

**Laine RF, Tosheva KL, Gustafsson N, Gray RDM, Almada P, Albrecht D, Risa GT, Hurtig F, Lindås A-C, Baum B, et al.** NanoJ: a high-performance open-source super-resolution microscopy toolbox. J Phys Appl Phys. 2019:**52**(16):163001. https://doi.org/10.1088/1361-6463/ab0261

**Lee H-C, Huang Y-P, Huang Y-W, Hu C-C, Lee C-W, Chang C-H, Lin N-S, and Hsu Y-H**. Voltage-dependent anion channel proteins associate with dynamic Bamboo mosaic virus-induced complexes. Plant Physiol. 2022:**188**(2):1061–1080. https://doi.org/10.1093/plphys/kiab519

**Li Y-H, Ke T-Y, Shih W-C, Liou R-F, and Wang C-W**. NbSOBIR1 Partitions Into Plasma Membrane Microdomains and Binds ER-Localized NbRLP1. Front Plant Sci. 2021:**12**. https://doi.org/10.3389/fpls.2021.721548

**Nelson BK, Cai X, and Nebenführ A**. A multicolored set of in vivo organelle markers for co-localization studies in Arabidopsis and other plants. Plant J. 2007:**51**(6):1126–1136. https://doi.org/10.1111/j.1365-313X.2007.03212.x

**Schindelin J, Arganda-Carreras I, Frise E, Kaynig V, Longair M, Pietzsch T, Preibisch S, Rueden C, Saalfeld S, Schmid B, et al.** Fiji: an open-source platform for biological-image analysis. Nat Methods. 2012:**9**(7):676–682. https://doi.org/10.1038/nmeth.2019

**Weber E, Engler C, Gruetzner R, Werner S, and Marillonnet S**. A modular cloning system for standardized assembly of multigene constructs. PloS One. 2011:**6**(2):e16765. https://doi.org/10.1371/journal.pone.0016765

**Wickham H**. Data Analysis. . In. ggplot2: Elegant Graphics for Data Analysis, H Wickham, ed. (Springer International Publishing: Cham), pp. 189–201. https://doi.org/10.1007/978-3-319-24277-4_9

**Win J, Kamoun S, and Jones AME**. Purification of Effector–Target Protein Complexes via Transient Expression in Nicotiana benthamiana. . In. Plant Immunity: Methods and Protocols, JM McDowell, ed. (Humana Press: Totowa, NJ), pp. 181–194. https://doi.org/10.1007/978-1-61737-998-7_15

**Wu C-H, Adachi H, De la Concepcion JC, Castells-Graells R, Nekrasov V, and Kamoun S**. NRC4 Gene Cluster Is Not Essential for Bacterial Flagellin-Triggered Immunity1 [OPEN]. Plant Physiol. 2020:**182**(1):455–459. https://doi.org/10.1104/pp.19.00859
